# Transcriptional pattern enriched for synaptic signaling is associated with shorter survival of patients with high-grade serous ovarian cancer

**DOI:** 10.1101/2024.10.21.619015

**Authors:** Arkajyoti Bhattacharya, Thijs S. Stutvoet, Mirela Perla, Stefan Loipfinger, Mathilde Jalving, Anna K.L. Reyners, Paola D. Vermeer, Ronny Drapkin, Marco de Bruyn, Elisabeth G.E. de Vries, Steven de Jong, Rudolf S.N. Fehrmann

## Abstract

**Background:** Bulk transcriptomic analyses of high-grade serous ovarian cancer (HGSOC) so far have not uncovered potential drug targets, possibly because subtle, disease-relevant transcriptional patterns are overshadowed by dominant, non-relevant ones. Our aim was to uncover disease-outcome-related patterns in HGSOC transcriptomes that may reveal novel drug targets.

**Method:** Using consensus-independent component analysis, we dissected 678 HGSOC transcriptomes of systemic therapy naïve patients—sourced from public repositories—into statistically independent transcriptional components (TCs). To enhance c-ICA’s robustness, we added 447 transcriptomes from non-serous histotypes, low-grade serous, and non- cancerous ovarian tissues. Cox regression and survival tree analysis were performed to determine the association between TC activity and overall survival (OS). Finally, we determined the activity of the OS-associated TCs in 11 publicly available spatially resolved ovarian cancer transcriptomes.

**Results:** We identified 374 TCs, capturing prominent and subtle transcriptional patterns linked to specific biological processes. Six TCs, age, and tumor stage stratified patients with HGSOC receiving platinum-based chemotherapy into ten distinct OS groups. Three TCs were linked to copy-number alterations affecting expression levels of genes involved in replication, apoptosis, proliferation, immune activity, and replication stress. Notably, the TC identifying patients with the shortest OS captured a novel transcriptional pattern linked to synaptic signaling, which was active in tumor regions within all spatially resolved transcriptomes.

**Conclusion:** The association between a synaptic signaling-related TC and OS supports the emerging role of neurons and their axons as cancer hallmark-inducing constituents of the tumor microenvironment. These constituents might offer a novel drug target for patients with HGSOC.

## Background

Epithelial ovarian cancer encompasses five primary histological subtypes, with high-grade serous ovarian carcinoma (HGSOC) constituting about 75% of all cases (1). The standard treatment for HGSOC diagnosed at stage IIB and beyond involves a combination of surgery and chemotherapy, primarily using platinum-based compounds and taxanes (2,3). While initial chemotherapy results in tumor response in most patients with HGSOC, there is a very high recurrence rate (4). The addition of poly-ADP ribose polymerase and vascular endothelial growth factor A inhibitors to chemotherapy for subsets of patients currently results in a 5-year disease-specific overall survival (OS) rate of approximately 45% for patients with HGSOC. This rate has hardly improved in the last three decades (5–8). Therefore, new insights into the complex biology underlying HGSOC are urgently needed to develop more effective treatment strategies.

Previous studies using bulk transcriptomes of patients with HGSOC have identified expression-based molecular subtypes. However, these subtypes did not provide insights that have translated into novel drug targets (9–11). A common limitation of such studies is their reliance on bulk transcriptomes, containing both tumor cells and tumor microenvironment (TME) components, thus reflecting the average transcriptional patterns of the combination of all biological processes present in the tumors. This averaging often masks subtle transcriptional patterns pivotal to understanding HGSOC biology, especially when these are overshadowed by dominant patterns from other less relevant (non-)biological processes (12). Consensus-independent component analysis (c-ICA) offers an alternative by decomposing such bulk transcriptomes into statistically independent transcriptional patterns (i.e., transcriptional components; TCs)(13,14). This approach reveals both dominant and subtle patterns and provides a measure of TC activity for each sample (15).

In the present study, our aim was to utilize c-ICA to dissect HGSOC transcriptomes to identify as many TCs associated with patient OS as possible, which could reveal potential novel drug targets.

## Methods

See Supplementary methods for the extended methods.

### Data acquisition

Raw microarray bulk transcriptomes and clinicopathological details for patients with HGSOC, low-grade serous ovarian cancer (LGSOC), non-serous ovarian cancer, and benign ovarian tissues were sourced from the Gene Expression Omnibus (GEO)(16). We exclusively utilized transcriptomes generated from primary tumor samples. Our analysis was confined to samples on the Affymetrix HG-U133 Plus 2.0 platform (GEO accession identifier: GPL570) and excluded cell line samples. The datasets were pre-processed, and quality controlled as previously described (17). Furthermore, for comprehensive analyses, we incorporated transcriptomes from five distinct resources: the Cancer Cell Line Encyclopedia (CCLE, n = 969), Genomics of Drug Sensitivity in Cancer (GDSC, n = 959), Gene Expression Omnibus (GEO, n = 13,810), and The Cancer Genome Atlas (TCGA, n = 8,150), and spatially resolved transcriptomes from 10xGenomics (16,18–20).

### Consensus-independent component analysis (c-ICA)

To preprocess the bulk transcriptome data, we applied a whitening transformation to prepare it for subsequent analysis. Consensus-ICA was conducted as described previously (Knapen et al., 2024). The output of an c-ICA includes two matrices: (i) transcriptional components (TCs) with gene weights, where each weight within the TC represents both the direction and magnitude of its effect on the expression levels of each gene, and (ii) a consensus mixing matrix (MM) with its coefficients representing the activity scores of TCs across samples.

### Survival analysis

To discern the relationship between TC activity and patient OS, a univariate Cox proportional hazards analysis was conducted on a select group of patients with available follow-up data (*n* = 541, Supplementary Table S1). In addition, a multivariate Cox proportional hazards analysis was carried out, including covariates such as age, stage, debulking status, and tumor grade. This latter analysis was based on a subset of patients with comprehensive clinicopathological data available (*n* = 373, Supplementary Table S1). We implemented a multivariate permutation framework encompassing 10,000 permutations to mitigate the risk of false discoveries. We established the acceptable false discovery rate (FDR) at 1%, maintaining an 80% confidence level, applicable for both the univariate and multivariate analyses.

### Survival tree analysis

We performed a survival tree analysis to delineate groups of patients with HGSOC treated with platinum-based chemotherapy based on distinct transcriptional and clinicopathological attributes. The analysis utilized activities of TCs associated with OS (either from univariate or multivariate survival analysis as mentioned in supplementary methods) in conjunction with relevant clinicopathological factors, such as age, tumor stage, debulking status, and grade, as potential classifiers. We divided patients into two subsets using every plausible cut-off point for each classifier and compared the resulting survival curves employing the log-rank statistic. Consequently, the division was based on the most significant classifier at its optimal cut-off based on the smallest p-value of the log-rank test mentioned above. This divisional process was successfully reiterated on the derived subsets until any of the following stipulated conditions was satisfied: *i*) the total patient count across both subsets fell below 50, *ii*) the collective number of uncensored events in both subsets was < 25, or *iii*) one of the subsets contained < 17 patients. To gauge the stability of our classifiers, we performed 20,000 iterations, randomly selecting 80% of the patient group in each iteration. The significance-based ranks of classifiers in these iterations were correlated with those from the primary survival tree.

### Associating the identified transcriptional components with biological processes

To discern the biological processes associated with the TCs, we adopted a multifaceted approach encompassing *i*) Transcriptional Adaptation to Copy Number Alterations (TACNA) profiling, targeting the identification of TCs that reflect the downstream implications of copy number alterations (CNAs) on gene expression levels(21); *ii*) Execution of gene set enrichment analysis (GSEA) for each TC, utilizing gene set collections (*n* = 16) from The Human Phenotype Ontology (The Monarch Initiative), the Mammalian Phenotypes (Mouse Genome Database), and the Molecular Signatures Database (MsigDB)(22,23); *iii*) The formation of co-functionality networks on the top and bottom genes of each TC, achieved using the GenetICA methodology, accessible via https://www.genetica-network.com(24). For clusters comprising ≥ 5 genes, the enrichment of the predicted functionality was quantified. This served as the foundation for determining the biological process associated with the TC being examined.

### Cross-study transcriptional component projection

To determine whether a biological process captured by an identified TC is also active in other cancer types and to investigate if it is more active in tumor cells or in the tumour microenvironment (TME), we collected raw expression profiles from multiple sources: the Cancer Cell Line Encyclopedia (CCLE, *n* = 969), Genomics of Drug Sensitivity in Cancer (GDSC, *n* = 959), Gene Expression Omnibus (GEO, *n* = 13,810), and The Cancer Genome Atlas (TCGA, *n* = 8,150)(16,18–20). While the CCLE and GDSC datasets comprise cell line profiles across many solid and hematologic malignancies, the GEO and TCGA datasets offer an extensive set of bulk transcriptomes derived from patient samples spanning 27 tumor types. We pre- processed the raw data as previously described (21). Next, we projected the TCs identified via c-ICA onto the cell line expression profiles from CCLE and GDSC and the patient-derived expression profiles from GEO and TCGA. This projection methodology has been described in more detail previously (21). To identify potential variations in the activity scores of the TCs, we compared the activity scores among cell lines and samples derived from patients within all four repositories. We used an absolute activity score threshold of 0.05 for each TC to pinpoint outlier cell lines and patient tumors with heightened activity.

### Determination of spatial transcriptomic profiles’ significant activity locations for individual transcriptional components

To further assess whether a biological process captured by an identified TC is more active in tumor cells or in the TME, we collected publicly available spatial resolved transcriptomic profiles of ovarian cancer samples. Eight were sourced from GEO (study ID GSE211956), and three were sourced from the public dataset repository of 10xGenomics (see supplementary methods for details) generated using the 10xGenomics Visium platform. The samples were from patients with HGSOC, serous papillary, and endometrioid ovarian cancer. Activity for each TC across every location within the spatial samples was ascertained through the cross-study projection methodology referred to in the previous method section (21). We incorporated a permutation-driven approach to discern the markedly active areas within the spatial samples for each TC. We derived a null distribution of activities for each TC-location pairing by performing 3,000 permutations and subsequent projections. The p-value of each observed TC activity quantifies the significance of the deviation of the TC activity at a given location from its baseline null distribution. After this, we visualized the z-transformed p-values using a heatmap, followed by obtaining colocalization scores for each combination of TCs in the spatial transcriptomic profiles for each ovarian cancer sample (25). This visualization aided in highlighting the areas with notable activity aligned against the stained representation of the tissue sample.

## Results

### An integrated data set containing 1,125 bulk transcriptomes from ovarian tissues

We curated 1,193 bulk transcriptomes from the GEO, including patients with HGSOC, low-grade serous ovarian cancer (LGSOC), non-serous ovarian cancer, and benign ovarian tissues (16). These were extracted from 32 distinct studies and represented the entire spectrum of ovarian cancer types, stages, and grades, and included 43 samples from non-malignant ovarian tissue. Pre-processing, which included removing duplicates and quality checks, culminated in a refined dataset of 1,125 samples (17). Supplementary Tables S1 and S2 provide detailed breakdowns of these samples, showcasing the comprehensive coverage of ovarian cancer types, stages, and grades within this dataset. The ovarian cancer dataset comprised bulk transcriptomes of patients with HGSOC (*n* = 678), other serous (*n* = 110), endometrioid (*n* = 110), and clear-cell ovarian cancer samples (*n* = 96). Additionally, for 541 patients, comprehensive survival data was available, as well as additional clinicopathologic information, including age, grade, stage, subtype, treatment schedule, and debulking status for 373 patients (Fig. 1).

**Figure 1.**
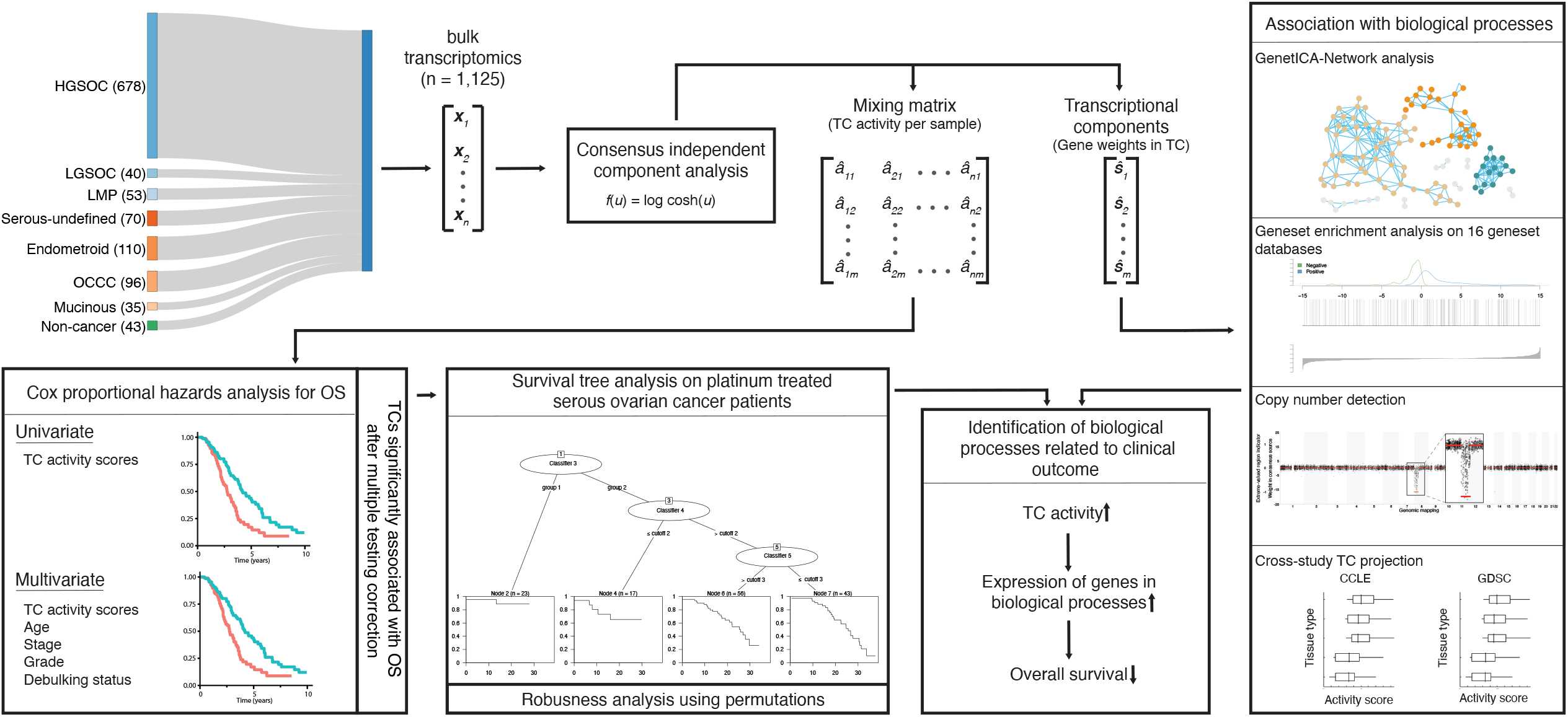
Workflow indicates the data acquisition and relations between the methods.

### Consensus-independent component analysis identifies 374 transcriptional components (TCs)

c-ICA on the 1,125 bulk transcriptomes revealed 374 independent TCs. Notably, 135 TCs captured the impact of copy number alterations on gene expression levels. Each TC displayed enrichment for at least one gene set from the 16 gene set collections, with an absolute Z-score of more than two. For example, the number of enriched gene sets from the Hallmark gene set collection in an individual TC ranged from zero to 28 enriched gene sets (interquartile range 3 – 7). The median top Z score for Hallmark gene sets was 3.21 (range 1.55 – 37.54, interquartile range 2.6 – 4.25). A comprehensive database, including all TCs and GSEA outcomes, has been made accessible at http://transcriptional-landscape-ovarian.opendatainscience.net.

The activities of 13 TCs were associated with patient overall survival (OS) in a univariate analysis, with an additional TC (TC166) identified in a multivariate analysis accounting for age, stage, debulking, and tumor grade. Combined, these 14 OS-associated TCs were enriched for gene sets associated with diverse biological processes and clinicopathological characteristics, with four TCs capturing the effects of copy number alterations on gene expression levels.

### The activities of six transcriptional components are associated with patient overall survival

For a selected subset of 541 patients—including HGSOC, LGSOC, and non-serous ovarian cancer—with available OS information (Supplementary Table S1), 13 TC activities displayed an association with OS univariately (false discovery rate 5%, confidence level 80% in permutation-based multiple testing framework Supplementary Table S3; Fig. 2). For patients with serous ovarian cancer, treated with platinum-based therapy (*n* = 301, Supplementary Table S1), lower activity of one additional TC (TC166) was associated with worse OS independent of age, stage, debulking, and tumor grade (Supplementary Table S4). Combined, these 14 OS-associated TCs were enriched for gene sets associated with diverse biological processes and clinicopathological characteristics. Four of these TCs captured the downstream effects of CNAs on gene expression levels (Fig. 2, Supplementary Fig. S1-S3). Survival tree analysis identified ten groups of patients with platinum-treated HGSOC based on the activity of six OS-associated TCs and the presence of two clinicopathological characteristics, namely age and stage (Fig. 3, Supplementary Table S5, median robustness statistic of survival tree = 0.52, interquartile range = 0.36 - 0.69). The survival tree demonstrated good classification power (concordance statistic = 0.72, standard error = 0.021). As expected, patients were divided into separate survival groups based on stage (1/2 vs. 3/4) and age (<53.7 vs. ≥53.7 years). The most significant difference in OS was observed between the cohorts with low and high TC121 activity (Supplementary Table S5). Patients with high TC121 tumor activity exhibited the shortest OS, also observed for the subset of patients with advanced-stage HGSOC (Supplementary Fig. S4, Supplementary Table S6). Supplementary Fig. S5 indicates that TC121 activity is highest in patients with HGSOC compared to other ovarian cancer subtypes. Notably, all subtypes contain subsets of samples with elevated TC121 activity. These robust associations with OS for TC121 in these two subsets of patients indicate the relevance of TC121, irrespective of stage.

**Figure 2.**
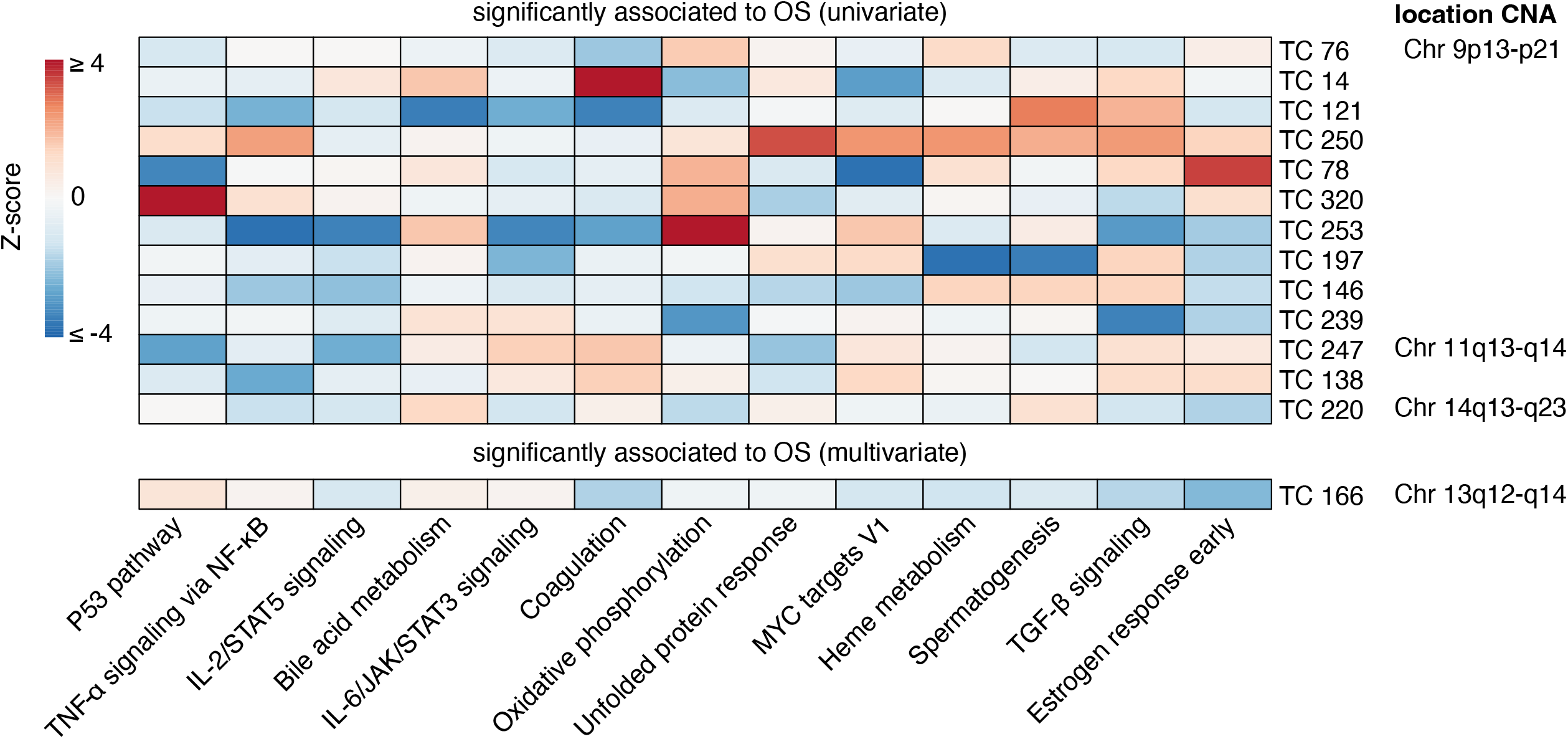
Enrichment heatmap of hallmark gene sets in transcriptional components associated with patient overall survival. Gene Set Enrichment Analysis for 14 TCs associated with OS identified through univariate or multivariate survival analyses are presented. Only Hallmark gene sets with significant enrichment (Bonferroni-corrected p-value) for at least one TC are shown. The heatmap displays Z-scores, which indicate the relative enrichment strength, with values truncated at a maximum of 4 for visualization purposes. The gene sets were clustered based on Pearson correlation using the Ward D2 method, providing insights into related biological processes captured by different TCs. In the right column, chromosomal locations of copy number alterations (CNAs) are shown, reflecting the downstream effects on gene expression that each TC captures. This integration of CNA information highlights the biological relevance of each TC and its contribution to gene expression variability and patient outcomes.

**Figure 3.**
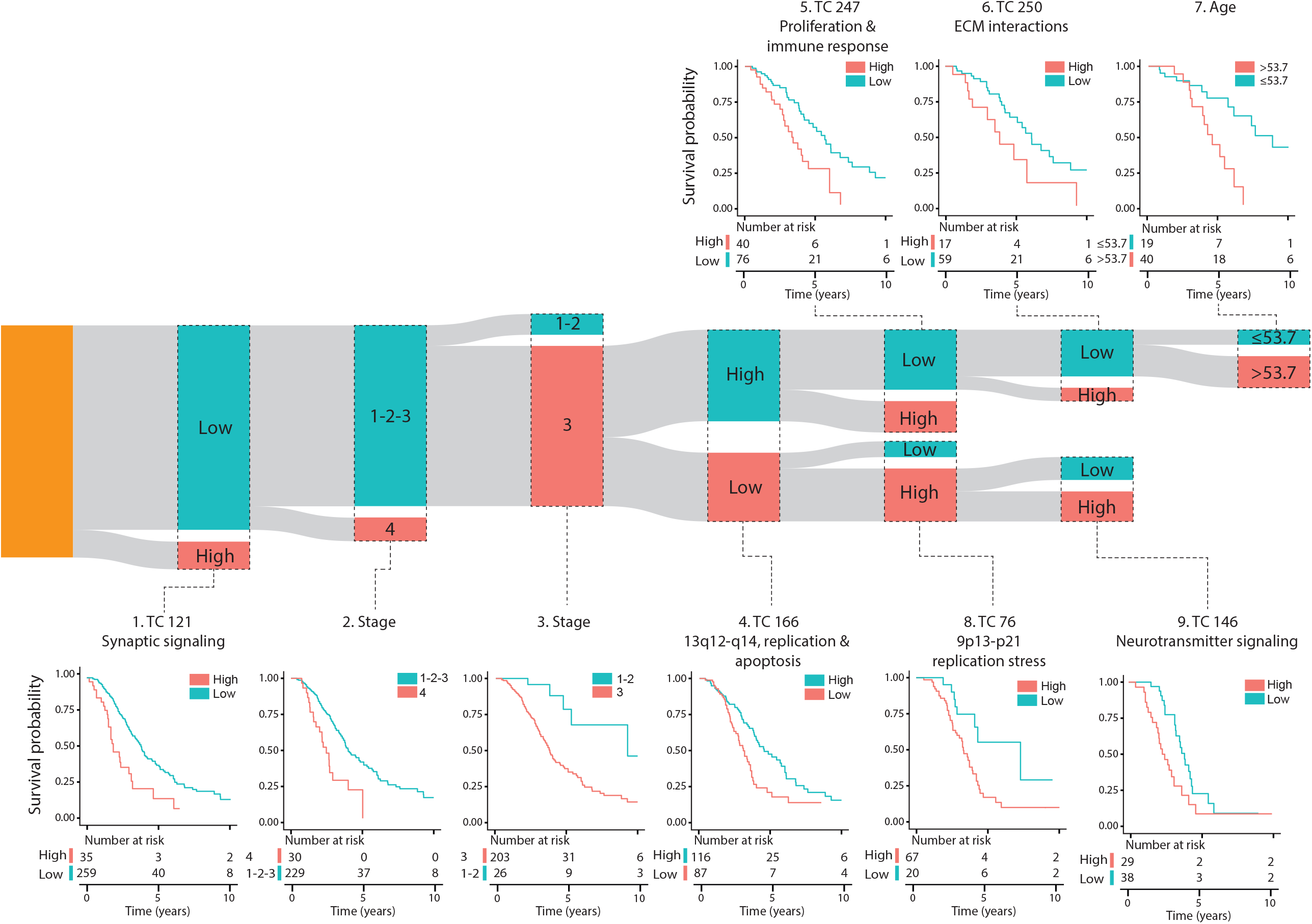
Survival tree analysis of patients with platinum-treated HGSOC defines survival cohorts with distinct clinicopathologic and biological characteristics. The results of survival tree analysis of 294 patients with high-grade serous ovarian cancer (HGSOC) treated with platinum-based chemotherapy are presented. The analysis utilized 14 transcriptional components (TCs) associated with overall survival (OS), along with other clinicopathologic factors, including age, tumor stage, grade, and debulking status. The resulting tree identified nine distinct survival cohorts, each represented as a bar in the Sankey diagram, where the bar height corresponds to the number of patients in each cohort. Kaplan-Meier survival curves with accompanying number-at-risk tables are shown for each cohort, with survival data censored at 10 years. The names of the survival cohorts were based on enriched biological processes in the TCs, as determined by the chromosomal location of genes captured by a TC, GSEA, and co-functionality analysis of the top genes. The p-values in the Kaplan-Meier plots were derived from log-rank tests comparing survival distributions between groups. Abbreviations: TC = transcriptional component, ECM = extracellular matrix.

### Distinct biological processes show enrichment in the transcriptional components associated with overall survival

Three of the six TCs associated with OS—TC166, TC247, and TC76—captured the effects of CNAs on the expression levels of genes mapping to chromosome regions 13q12-q14, 11q13-q14, and 9p13-p21, respectively (Supplementary Fig. S6, Supplementary Table S7) (6). The higher activity of TC166 was associated with better OS, whereas the higher activities of TC121, TC247, TC250, TC76, and TC146, were associated with worse OS. Among the 14 OS-associated TCs, only TC166 showed a significant association with OS in an independent cohort of patients with ovarian clear cell carcinoma (Bonferroni corrected p-value < 0.05; see supplementary methods and Supplementary Table S8) (26). The top genes from TC166 were enriched for genes involved in replication and apoptosis. The chromosomal region 13q12- q14 linked to the TC166, contains the tumor suppressor genes retinoblastoma 1 (*RB1*) and Breast Cancer Type 2 Susceptibility Protein (*BRCA2*). Loss of heterozygosity of this chromosomal region is frequently observed in both sporadic and hereditary serous ovarian cancers (27,28). The top genes from TC247 were enriched for genes involved in proliferation and immune cell activation, TC76 in replication stress, TC250 in extracellular matrix (ECM) interactions, and TC146 in neurotransmitter signaling.

Intriguingly, the top 100 genes in TC121 revealed a co-functional cluster enriched for genes involved in synaptic signaling, with the corresponding proteins reported to localize to the synaptic membrane of neurons (Fig. 4). Among these were pre-synaptic protein neurexin-1 (*NRXN1*) and its post-synaptic ligand leucine-rich repeat transmembrane protein 2 (*LRRTM2*), which regulates excitatory synapse formation (top 20 genes are described in Supplementary Table S9, for more details: http://transcriptional-landscape-ovarian.opendatainscience.net) (29,30). Furthermore, this co-functional cluster included neuron-specific synaptic structure proteins, neurofilament light, and medium chain. Moreover, genes encoding for potassium ion channel proteins integral to membrane repolarization during synapse signal transduction carried high weights in TC121. These genes included *KCNC1, KCNN2*, and *KCNIP1*(31–33). Several genes in TC121 encoded proteins related to glutamate receptor signaling, including *GRIN2C* and *SLC7A10* (34). In line with this proposed function, high activity of TC121 was observed in neuroblastoma cell lines but not in ovarian or central nervous system cancer cell lines in the GDSC and CCLE datasets (Fig. 5A, Supplementary Fig. S7, S8). In the GEO and TCGA datasets, high activity of TC121 was observed in glioblastoma multiforme and lower-grade glioma but not in ovarian cancer patient samples (Fig. 5B).

**Figure 4.**
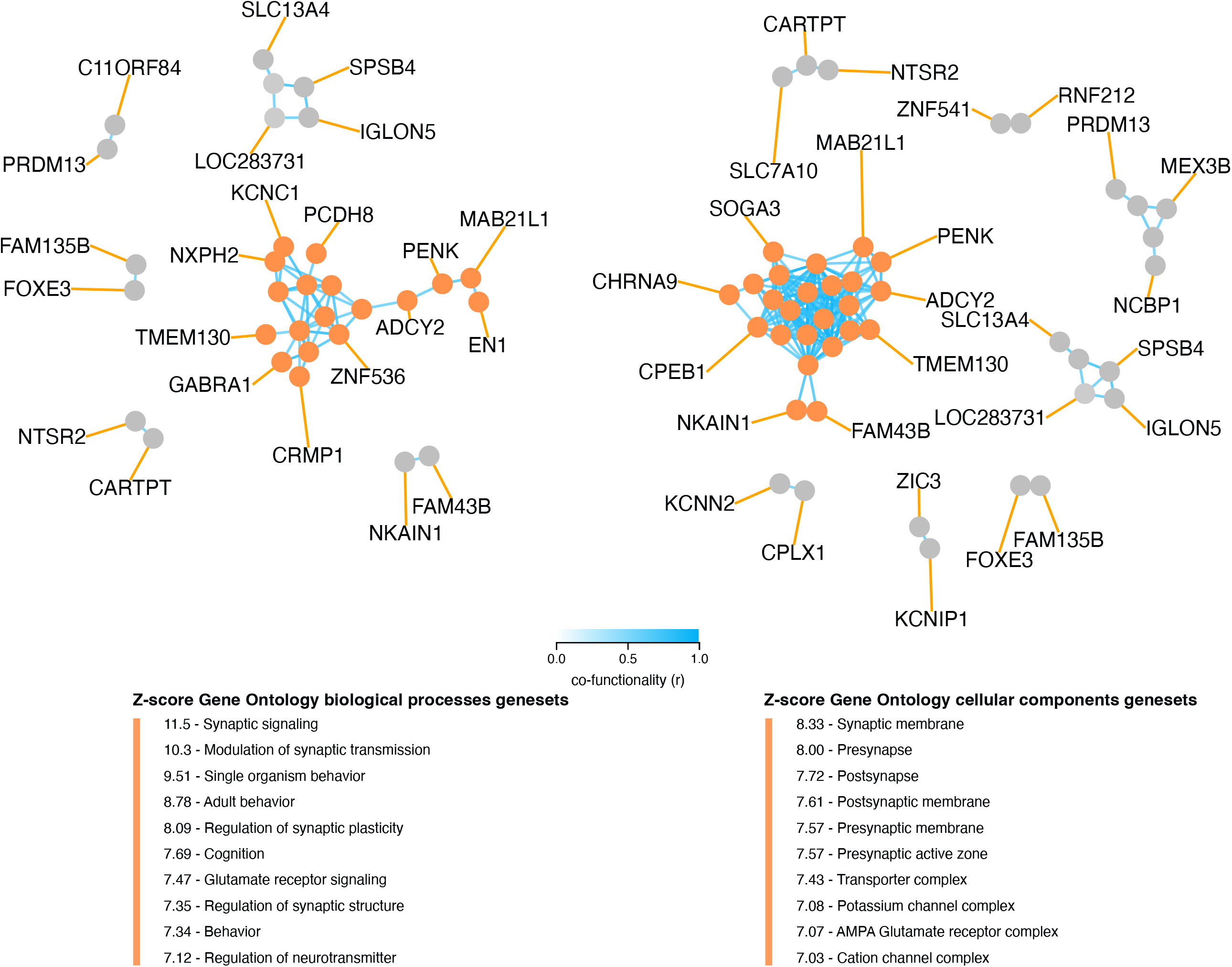
Co-functionality network of top 100 absolute weighted genes in TC121. Co-functionality network for the top 100 genes with the highest absolute weights in TC121 is presented. Genes were clustered based on predicted co-functionality (r > 0.7) across datasets, with clusters identified using both Gene Ontology (GO) Biological Processes and Cellular Components databases. One primary cluster, containing more than five genes, exhibited strong enrichment for synaptic signaling in the GO Biological Processes database and for synaptic membranes in the GO Cellular Components database. This highlights the biological specificity of TC121 in regulating gene expression linked to synaptic functions.

**Figure 5.**
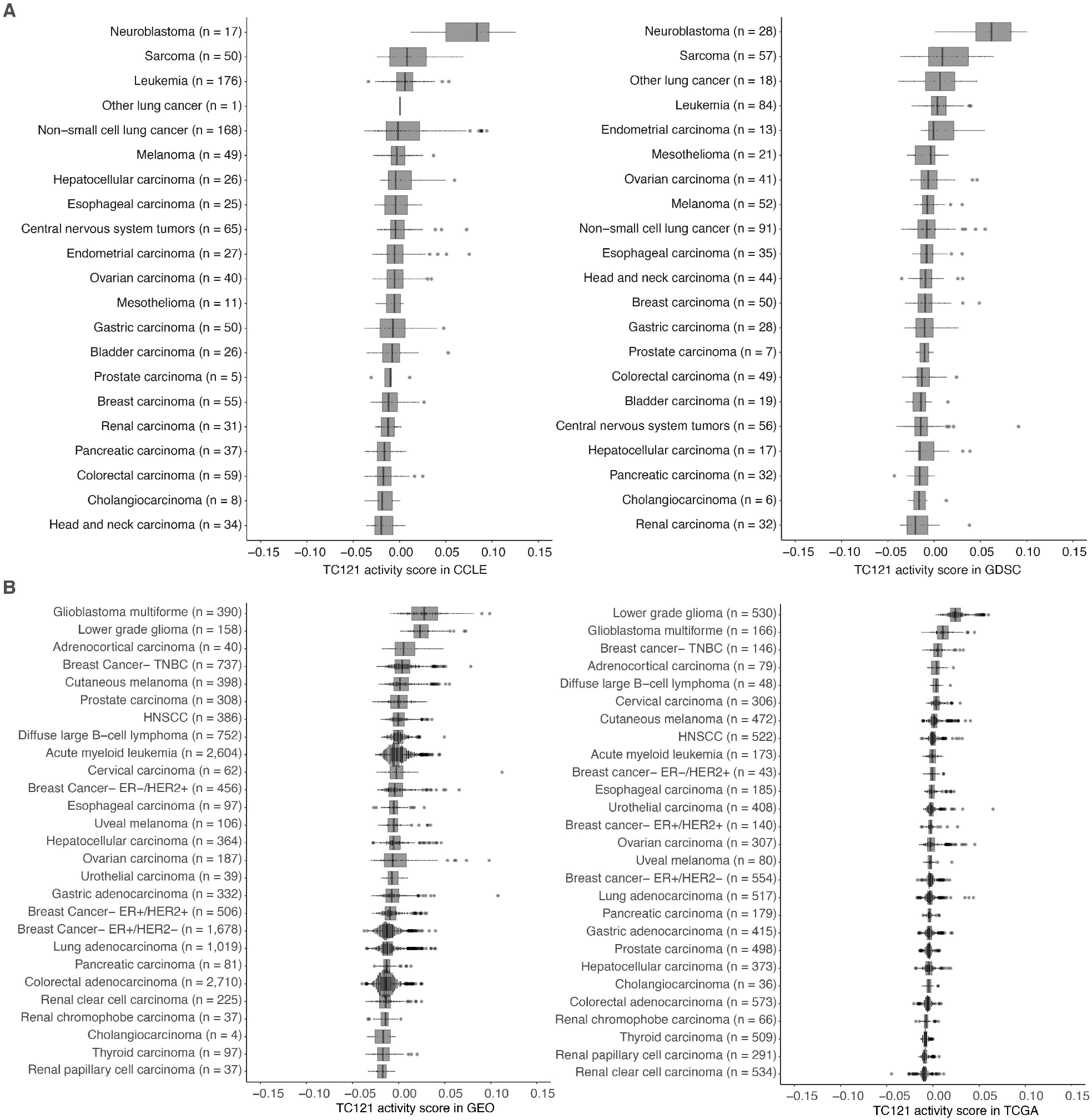
The activity of TC121 in bulk transcriptomes of CCLE, GDSC cell lines, and GEO and TCGA patient-derived samples. **A.** Cross-study TC projection of TC121 on CCLE and GDSC cell lines. The boxplots display the activity scores of TC121 in different tissue types, which are ordered based on their corresponding median activity scores. **B**. Cross-study TC projection of TC121 on GEO and TCGA bulk transcriptomes resulted in the activity scores presented in the boxplots. Cancer types were ordered based on corresponding medians of TC121 activity scores. Abbreviations: TC = transcriptional component.

### Distinct cluster of patients from TCGA overlap with elevated activity of TC121

To explore if pre-existing classifications of patients with ovarian cancer correspond to the contrasting activities of the TCs, we investigated the classification provided by TCGA. TCGA identified four optimal clusters describing the patients with ovarian cancer using transcriptional profiles.(9) To explore associations between these clusters and TC activity, we performed a Kruskal-Wallis test using TCGA sample data. Supplementary Fig. S9 highlights the associations between each cluster set and the TCs, represented by log-transformed p-values. A significant association between a TC and a cluster set indicates that at least one cluster within the cluster set exhibited significantly different activity scores for the corresponding TC compared to the other clusters. Notably, samples with high TC121 activity were not captured by any of the clusters of the four-cluster set. Interestingly, the eight-cluster set predefined by TCGA was able to identify a cluster that corresponded to samples with elevated TC121 and TC146 activity. This finding suggests that while TCGA’s analysis identified this patient group based on transcriptional profiles, it didn’t characterize them further.

### Distinct spatial and single cell transcriptional profiles with high activity of OS-associated TCs

Cross-study TC projection onto spatial transcriptomic profiles from 11 ovarian cancer samples revealed that TC121 was highly active in profiles from the tumor region of the 11 ovarian cancer samples (Fig. 6A; Supplementary Fig. S10). Additionally, TC121 showed markedly higher activity in the transcriptional profiles of a subset of the unannotated single cells from HGSOC patients (study ID GSE158722; see supplementary methods and Supplementary Fig. S11).(35) This finding suggests that some of these unannotated cells could be neurons. Furthermore, the unannotated single cell transcriptional profiles showed contrasting activity scores of different OS-associated TCs (supplementary Fig. S11). These contrasting activities indicate that these TCs could provide insights into the biology of previously uncharacterized cell types. Distinct regions with high activity of the copy number TCs (TC166, TC247, and TC76) in the HGSOC sample overlapped with the region containing cancer cells, as expected. TC250, enriched for extracellular matrix interactions, was also active in the stromal region. The strongest inverse colocalization (colocalization score -2.43) was observed between the activity scores of TC146, enriched for neurotransmitter signaling, and TC76, captured the effect of copy number alterations at chromosome 9p13-p21, at the serous ovarian cancer sample (Fig. 6B, Supplementary Table S10).

**Figure 6.**
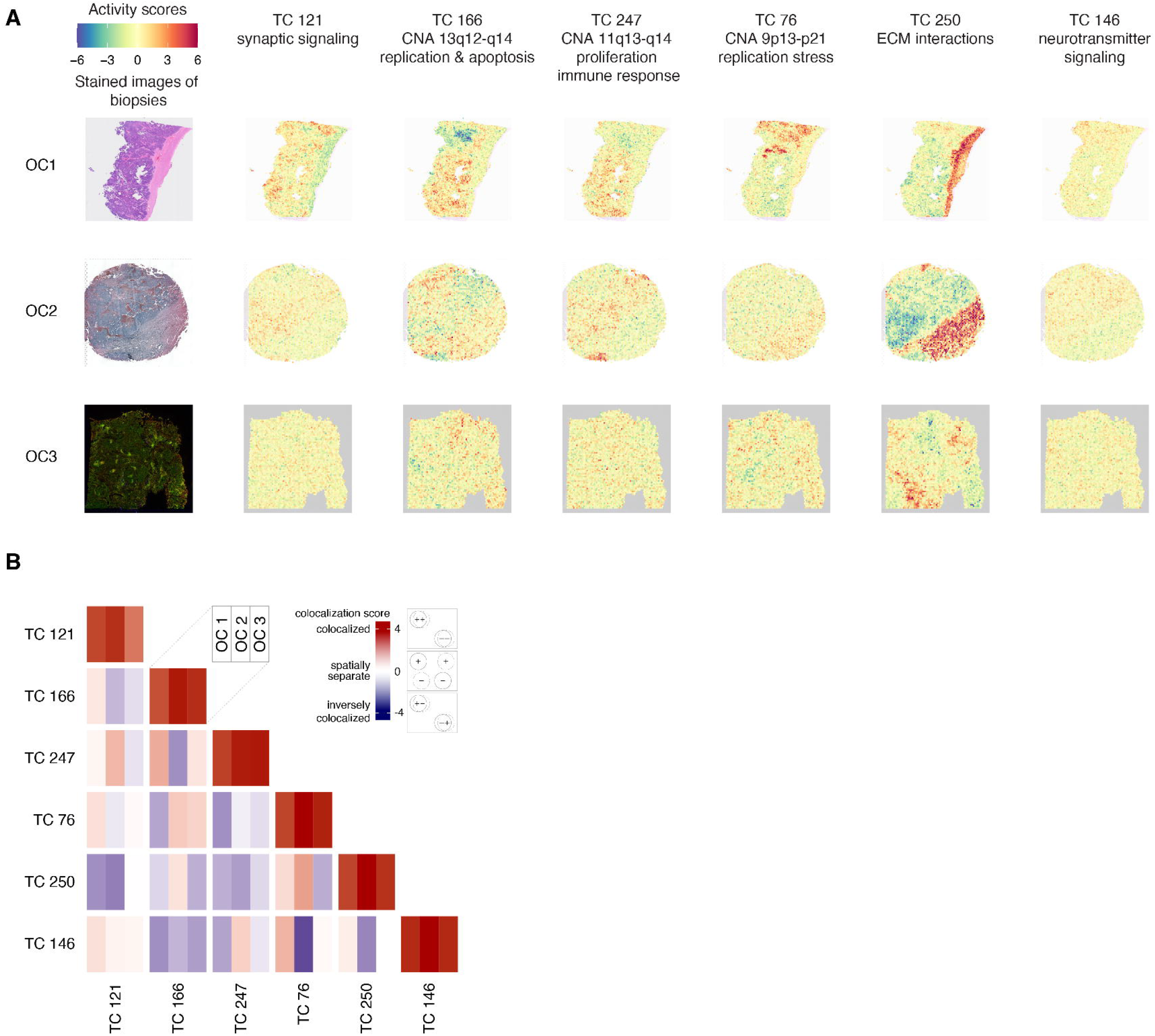
Spatial transcriptomic profiles in ovarian cancer samples. **A**. We employed a permutation-based approach to pinpoint the areas of significant TC activity in spatial transcriptomic profiles. We ran 5,000 permutations for each TC-profile combination, yielding a p-value that indicates the extent to which the TC activity in the corresponding profile differs from what would be expected by chance (the null distribution). We then transformed these p-values into logarithmic values and represented them using a heatmap. Heatmaps of activity scores of the TCs are presented in individual rows for the HGSOC, serous papillary, and endometrioid adenocarcinoma of ovary samples. The first column represents the stained images of the samples. The second to seventh columns show heatmaps corresponding to the mentioned TCs. **B**. The heatmap illustrates the colocalization between two TC activities on spatial transcriptomic profiles from ovarian cancer samples. For each cell, the colocalization scores of the TCs at each of the three spatial transcriptomics samples OC 1, OC 2, and OC 3 are arranged in columns. A colocalization score of 4 between two TCs (red) indicates that the positively (+) and negatively (−) active regions of both TCs are perfectly colocalized. Conversely, a colocalization score of -4 between two TCs (blue) also indicates colocalization. Still, with inverse activity, i.e., the positively active regions of one TC are colocalized with the negatively active regions of the other TC or vice versa. A colocalization score close to 0 between two TCs (white) indicates that the activities of two TCs are spatially separated. The dashed and solid circle in the panel on the right side of the color bar represents two different TCs. Abbreviations: TC = transcriptional component.

## Discussion

In this study, we identified 374 TCs, each enriched for gene sets representing various biological processes in HGSOC samples. Six could stratify patients with HGSOC who had received platinum-based treatment into ten distinct OS groups.

The most significant TC in the survival tree analysis, TC121, captured a clinically relevant subtle transcriptional pattern linked to synaptic signaling not previously recognized in HGSOC. In the survival tree, TC121 identified 12% of the HGSOC patients with the shortest OS and, based on spatially resolved transcriptomic analyzed samples, is active in tumor regions. This observation supports the emerging role of neurons and neuronal projections as cancer hallmark-inducing constituents of the TME (36–38).

Further investigation on whether the activity of TC121 originated from tumor cells or in the TME revealed that the TC121 signal is coming from cells within the TME. The high activity of TC121 in low-grade glioma and glioblastoma multiforme patient samples (Fig. 5B) is in agreement with the presence of neurons in large numbers within the TME of gliomas, where they form functional synapses with tumor cells(39,40). Moreover, TC121 activity was lower in non-brain cancers, such as ovarian cancers, which contain fewer neurons and synapses in the TME compared to brain cancers. We expected TC121 activity to be low in the bulk transcriptomes of all cell lines, since they lack TME. TC121 activity in most cell lines, which includes glioblastoma and ovarian cancer cell lines, was indeed low. Neuroblastoma cell lines, however, exhibited high TC121 activity, which is likely due to retained synaptic formation capacity originating from neuroblast cells (41,42). Lastly, TC121’s high activity observed in small, scattered regions within the tumor of spatially resolved transcriptomic ovarian cancer samples also supports TC121’s role in the TME.

TC121’s significant association with OS underscores the potential significance of synaptic signaling in HGSOC biology. Yet, the neuronal subtype and the molecular mechanisms associated with TC121 remain to be elucidated. A study in human ovarian cancer-bearing mice demonstrated that sympathetic innervation in HGSOC involves adrenergic signaling: norepinephrine released by sympathetic neurons binds to beta-adrenergic receptors on the cancer cells (43–45). This binding triggers the tumor cells to release brain-derived neurotrophic factor (BDNF), which enhances cancer innervation via activation of host neurotrophic receptor tyrosine kinase B receptors (NTRK2), thereby establishing a feed-forward loop of sustained signaling. BDNF and the nerve marker neurofilament protein expression were examined in 108 human ovarian tumors (41). This study revealed that increased intratumoral nerve presence strongly correlates with elevated BDNF and norepinephrine levels, advanced tumor stage, and shorter OS in patients with ovarian cancer. This interaction can be targeted with pan-TRK inhibitors such as entrectinib and larotrectinib. Both drugs are showing promising results in multiple phase II trials, including ovarian cancer and breast cancer patients. Furthermore, a TRKB-specific inhibitor was developed (ANA-12), but has not been subjected to any clinical trials in cancer so far (46–49). Our analysis indicated that *BDNF* is a prominent gene (with an absolute weight > 3) in 10 TCs but not in TC121, suggesting that TC121 may indicate a distinct process unrelated to BDNF.

The significance of sensory innervation in HGSOC was evidenced by the co- localization of TRPV1, a marker for sensory neurons, and β-III tubulin, a general neuronal marker, in immunofluorescent staining of histological sections from 75 patients (50). Additionally, a murine model study employing neural tracing identified sensory neurons originating from local dorsal root ganglia and jugular–nodose ganglia, with axons extending into the TME (50). A transgenic murine model lacking nociceptors demonstrated that this specific subtype of sensory neurons was involved in tumor progression (51). Another study showed that reducing the release of Calcitonin gene-related peptide from tumor-innervating nociceptors could be a strategy to alleviate this effect of nociceptors by improving anti-tumour immunity of cytotoxic CD8+ T cells in a melanoma model bearing mice(52). This indicates that the signal from TC121 may represent an indirect influence on tumor cells via interactions with immune cells and the promotion of an immune suppressive TME. Furthermore, in cell lines derived from Trp53^−/−^ Pten^−/−^ murine HGSOC, the influence of nociceptors was characterized by the release of substance P (SP), their primary neuropeptide. SP is an alternative splicing product of the preprotachykinin A gene (*TAC1*) and binds to the receptor neurokinin 1 (NK1R), encoded by the *TACR1* gene. NK1R expression was confirmed in murine HGSOC cell line, and SP enhanced cellular proliferation in NK1R-positive murine HGSOC cancer cells *in vitro* (51). Our analysis identified *TAC1* and *TACR1* as prominent genes in 15 and 2 TCs, respectively, yet not in TC121, and none of these TCs were associated with patient survival. Currently, there are no drugs specifically targeting tumor innervation in (ovarian) cancer (53). Interestingly, the NK1R antagonist aprepitant effectively inhibited the metastasis-promoting effects of neural substance P in human breast and mammary cancer bearing mice (54), demonstrating the feasibility of such an approach.

Strategies to disrupt neuronal signaling and neurotransmitter release in neurons target key elements of excitatory neurotransmission, such as calcium flux and vesicle formation. Drugs like ifenprodil and lamotrigine, commonly used to treat neuronal disorders, block glutamate release and subsequent neuronal signaling. Additionally, the vesicular monoamine transporter (VMAT) inhibitor reserpine prevents synaptic vesicle formation (55,56). In vitro studies with HGSOC cell lines have demonstrated that ifenprodil significantly inhibits tumour proliferation, while reserpine induces apoptosis in cancer cells (57,58). These approaches hold promise for inhibiting neuronal signaling and interactions in the TME. Therefore, it is essential that the mechanisms driving this nerve growth, the specifics of how nerves within the TME interact with ovarian cancer cells, and how they impact the survival of patients with HGSOC are further elucidated.

Altogether, the present study uncovered a clinically relevant TC linked to synaptic signaling not previously identified in HGSOC. This TC may represent a novel cancer cell-extrinsic mechanism within the TME, illustrating how cancer cells and nerve cells interact to promote enhanced proliferation. A deeper understanding of the molecular aspects of tumor innervation could pave the way for novel drug targets for patients with HGSOC.

## Data availability

Microarray expression data was collected from three public data repositories: Gene Expression Omnibus with accession number GPL570 (generated with Affymetrix HG-U133 Plus 2.0), CCLE (generated with Affymetrix HG-U133 Plus 2.0, file CCLE_Expression.Arrays_2013-03-18.tar.gz) available at https://portals.broadinstitute.org/ccle/data and GDSC (generated with Affymetrix HG-U219) available at https://www.ebi.ac.uk/arrayexpress/experiments/E-MTAB-3610/. Pre-processed and normalized RNA-seq data was collected from TCGA using the Broad GDAC Firehose portal (https://gdac.broadinstitute.org/). Spatially resolved samples were sourced from 10xGenomics and GEO. The datasets generated during and/or analyzed during the current study are available in the website: http://transcriptional-landscape-ovarian.opendatainscience.net. The accession numbers of the studies included in this analysis are mentioned in Supplementary Table S2.

## Code Availability

The complete set of codes utilized in this study is available at the github repository: https://github.com/arkajyotibhattacharya/TranscriptionalLandscapeOvarianCancer.

## Author contribution

A. Bhattacharya: Data curation, methodology, data analysis, data interpretation, writing. T. S. Stutvoet: Data curation, methodology, data analysis, data interpretation, writing. M. Perla: Data interpretation, writing. S. Loipfinger: Data analysis, data interpretation, writing. M. Jalving: Data interpretation, writing. A. K. L. Reyners: Data interpretation, writing. P. D. Vermeer: Data interpretation, writing. R. Drapkin: Data interpretation, writing. M. de Bruyn: Data interpretation, writing. E. G. E. de Vries: Data interpretation, writing. S. de Jong: Conceptualization, data interpretation, writing. R. S. N. Fehrmann: Conceptualization, data curation, methodology, data analysis, data interpretation, writing. All were involved in the final decision to submit the manuscript.

## Figure legends

**Supplementary Figure S1**. Association of OS-related TCs with clinicopathologic parameters. The association between clinicopathologic parameters and the activity of the OS-associated 14 TCs was determined in the complete set of 1,125 samples. Pearson correlation was used to calculate the association of each clinicopathologic parameter with each TC. The TCs were then ranked based on their association with OS. Abbreviations: TC = transcriptional component, OS = overall survival.

**Supplementary Figure S2**. Enrichment heatmap for the KEGG gene set collection in OS- related TCs. GSEA results of 14 TCs associated with OS in univariate or multivariate survival analysis are presented, including KEGG gene sets that were included if the enrichment for at least one TC passed the Bonferroni threshold for multiple testing correction. The gene sets were clustered using Pearson correlation and Ward D2, and the heatmap colors were based on Z-scores, truncated at a value of four. The right column shows the chromosomal location of a copy number alteration that the TC captures the downstream effects on gene expression levels. Abbreviations: TC = transcriptional component, OS = overall survival.

**Supplementary Figure S3**. Enrichment heatmap for the REACTOME gene set collection in OS- related TCs. GSEA results of 14 TCs associated with OS in univariate or multivariate survival analysis is presented, including REACTOME gene sets that were included if the enrichment for at least one TC passed the Bonferroni threshold for multiple testing correction. The gene sets were clustered using Pearson correlation and Ward D2, and the heatmap colors were based on Z-scores, truncated at a value of four. The right column shows the chromosomal location of a copy number alteration that the TC captures the downstream effects on gene expression levels. Abbreviations: TC = transcriptional component.

**Supplementary Figure S4**. Survival tree analysis of patients with advanced-stage, HGSOC defines survival cohorts with distinct clinicopathologic and biological characteristics. Survival tree analysis of 265 patients with advanced-stage, platinum-treated HGSOC using 14 OS- associated TCs and other classifiers such as age, tumor stage, grade, and debulking status. The analysis resulted in nine survival cohorts, and the height of the bar in the Sankey diagram represents the number of patients in each cohort. The Kaplan-Meier plots and number-at-risk tables are presented with survival data censored at 10 years. The names of the survival cohorts were based on enriched biological processes in the TCs, as determined by the chromosomal location of genes captured by a TC, GSEA, and co-functionality analysis of the top genes. The p-values in each panel show the p-value from the corresponding log-rank test between the two groups. Abbreviations: TC = transcriptional component, ECM = extracellular matrix.

**Supplementary Figure S5**. The activity of TC121 in bulk transcriptomes of patients with different subtypes of ovarian cancer. The boxplots display the activity scores of TC121 in different cancer subtypes, which are ordered based on their corresponding median activity scores.

**Supplementary Figure S6**. Three OS-associated TCs capture the transcriptional effect of copy number alterations. For each OS-related TC, the weight of each gene was plotted on its genomic location. Abbreviations: TC = transcriptional component.

**Supplementary Figure S7**. Activity of OS-associated TCs in bulk transcriptomes of CCLE cell lines. The proportion of samples with outlier activity scores in individual tissue types for each of the 14 OS-associated TCs was obtained using an absolute cut-off value of 0.05. TCs were ordered based on their association with OS. Abbreviations: TC = transcriptional component.

**Supplementary Figure S8**. The activity of OS-associated TCs in bulk transcriptomes of GDSC cell lines. The proportion of samples with outlier activity scores in individual tissue types for each of the 14 OS-associated TCs was obtained using an absolute cut-off value of 0.05. TCs were ordered based on their association with OS. Abbreviations: TC = transcriptional component.

**Supplementary Figure S9**. Association heatmap for the TCGA cluster-sets with the activity scores of OS-related TCs. This heatmap highlights the associations between each cluster set and the TCs, represented by log-transformed p-values from corresponding Kruskal-Wallis test. A significant association between a TC and a cluster set indicates that at least one cluster within the cluster set exhibited significantly different activity scores for the corresponding TC compared to the other clusters. The heatmap colors were based on log-transformed p-values. Abbreviations: TC = transcriptional component. OS = overall survival.

**Supplementary Figure S10**. Spatial transcriptomic profiles in eight ovarian cancer samples. **A**. We employed a permutation-based approach to pinpoint the areas of significant TC activity in spatial transcriptomic profiles. We ran 5,000 permutations for each TC-profile combination, yielding a p-value that indicates the extent to which the TC activity in the corresponding profile differs from what would be expected by chance (the null distribution). We then transformed these p-values into logarithmic values and represented them using a heatmap. Heatmaps of activity scores of the TCs are presented in individual rows for the HGSOC samples. The first column represents the stained images of the samples. The second to seventh columns show heatmaps corresponding to the mentioned TCs. **B**. The heatmap illustrates the colocalization between two TC activities on spatial transcriptomic profiles from ovarian cancer samples. For each cell, the colocalization scores of the TCs at each of the eight spatial transcriptomics samples are arranged in columns. A colocalization score of 4 between two TCs (red) indicates that the positively (+) and negatively (−) active regions of both TCs are perfectly colocalized. Conversely, a colocalization score of -4 between two TCs (blue) also indicates colocalization. Still, with inverse activity, i.e., the positively active regions of one TC are colocalized with the negatively active regions of the other TC or vice versa. A colocalization score close to 0 between two TCs (white) indicates that the activities of two TCs are spatially separated. The dashed and solid circle in the panel on the right side of the color bar represents two different TCs. Abbreviations: TC = transcriptional component.

**Supplementary Figure 11**. The activity scores of OS-associated TCs in different single cell types from patients with HGSOC. Abbreviations: TC = transcriptional component. OS = overall survival. HGSOC = high-grade serous ovarian cancer.

## Supplementary table legends

**Supplementary Table S1**. Patient characteristics. Abbreviations: NA = not applicable, HGSOC = high-grade serous ovarian cancer, LGSOC = low-grade serous ovarian cancer, LMP = low malignant potential, OCCC = ovarian clear cell cancer.

**Supplementary Table S2**. Overview of the number of samples from each GEO series and the corresponding study. Abbreviations: PMID = PubMed identification number.

**Supplementary Table S3**. Multivariate permutation framework results from the univariate survival analysis. Abbreviations: TC = transcriptional component. CI = confidence interval

**Supplementary Table S4**. Permutation framework results from the multivariate survival analysis. Abbreviations: TC = transcriptional component. CI = confidence interval

**Supplementary Table S5**. Clinicopathological parameters per survival tree node in platinum-treated HGSOC. Abbreviations: HGSOC = high-grade serous ovarian cancer, TC = transcriptional component, ECM = extracellular matrix.

**Supplementary Table S6**. Clinicopathological parameters per survival tree node in advanced-stage platinum-treated HGSOC. Abbreviations: HGSOC = high-grade serous ovarian cancer, TC = transcriptional component, ECM = extracellular matrix.

**Supplementary Table S7**. GenetICA gene network analysis results. Abbreviations: TC = transcriptional component.

**Supplementary Table S8**. Results from the univariate survival analysis on patients with ovarian clear cell carcinoma. Abbreviations: TC = transcriptional component. CI = confidence interval.

**Supplementary Table S9**. Expression and function of the top 20 genes in TC121, according to literature. Abbreviations: TC = transcriptional component.

**Supplementary Table S10**. Colocalization scores of every two TC combinations in spatial transcriptomic profiles from three ovarian cancer samples. Abbreviations: TC = transcriptional component.

## References

1. Lheureux S, Gourley C, Vergote I, Oza AM. Epithelial ovarian cancer. Lancet 2019;393:1240–53.

2. NCCN clinical practice guidelines in oncology - ovarian cancer, including fallopian tube cancer and primary peritoneal cancer. November 26, 2019.

3. Wright AA, Bohlke K, Armstrong DK, Bookman MA, Cliby WA, Coleman RL, et al. Neoadjuvant chemotherapy for newly diagnosed, advanced ovarian cancer: Society of Gynecologic Oncology and American Society of Clinical Oncology Clinical Practice Guideline. J Clin Oncol 2016;34:3460–73.

4. Corrado G, Salutari V, Palluzzi E, Distefano MG, Scambia G, Ferrandina G. Optimizing treatment in recurrent epithelial ovarian cancer. Expert Rev Anticancer Ther 2017;17:1147–58.

5. Cibula D, Balmaña J. PARP inhibitors in ovarian cancer. Ann Oncol 2016;27:i40–44.

6. Wang H, Xu T, Zheng L, Li G. Angiogenesis inhibitors for the treatment of ovarian cancer. Int J Gynecol Cancer 2018;28:903–14.

7. Wu SG, Wang J, Sun JY, He ZY, Zhang WW, Zhou J. Real-world impact of survival by period of diagnosis in epithelial ovarian cancer between 1990 and 2014. Front Oncol 2019;9:639.

8. Tewari KS, Burger RA, Enserro D, Norquist BM, Swisher EM, Brady MF, et al. Final overall survival of a randomized trial of Bevacizumab for primary treatment of ovarian cancer. J Clin Oncol 2019;37:2317–28.

9. Bell D, Berchuck A, Birrer M, Chien J, Cramer DW, Dao F, et al. Integrated genomic analyses of ovarian carcinoma. Nature 2011;474:609–15.

10. Tothill RW, Tinker AV, George J, Brown R, Fox SB, Lade S, et al. Novel molecular subtypes of serous and endometrioid ovarian cancer linked to clinical outcome. Clin Cancer Res 2008;14:5198–208.

11. Verhaak RGW, Tamayo P, Yang JY, Hubbard D, Zhang H, Creighton CJ, et al. Prognostically relevant gene signatures of high-grade serous ovarian carcinoma. J Clin Investig 2013;123:517–25.

12. Chen C, Grennan K, Badner J, Zhang D, Gershon E, Jin L, et al. Removing batch effects in analysis of expression microarray data: an evaluation of six batch adjustment methods. PLoS One 2011;6:e17238.

13. Kong W, Vanderburg CR, Gunshin H, Rogers JT, Huang X. A review of independent component analysis application to microarray gene expression data. BioTechniques 2008;45:501–20.

14. Chiappetta P, Roubaud MC, Torrsani B. Blind source separation and the analysis of microarray data. J Comput Biol 2004;11:1090–109.

15. Biton A, Bernard-Pierrot I, Lou Y, Krucker C, Chapeaublanc E, Rubio-Pérez C, et al. Independent component analysis uncovers the landscape of the bladder tumor transcriptome and reveals insights into luminal and basal subtypes. Cell Rep 2014;9:1235–45.

16. Clough E, Barrett T. Statistical genomics, methods and protocols. Methods Mol Biol 2016;1418:93–110.

17. Fehrmann RSN, Karjalainen JM, Krajewska M, Westra H-J, Maloney D, Simeonov A, et al. Gene expression analysis identifies global gene dosage sensitivity in cancer. Nat Genet 2015;47:115–25.

18. Barretina J, Caponigro G, Stransky N, Venkatesan K, Margolin AA, Kim S, et al. The cancer cell line encyclopedia enables predictive modelling of anticancer drug sensitivity. Nature 2012;483:603–7.

19. Yang W, Soares J, Greninger P, Edelman EJ, Lightfoot H, Forbes S, et al. Genomics of drug sensitivity in Cancer (GDSC): a resource for therapeutic biomarker discovery in cancer cells. Nucleic Acids Res 2013;41:D955–61.

20. Barrett T, Wilhite SE, Ledoux P, Evangelista C, Kim IF, Tomashevsky M, et al. NCBI GEO: archive for functional genomics data sets—update. Nucleic Acids Res 2013;41:D991–5.

21. Bhattacharya A, Bense RD, Urzúa-Traslaviña CG, de Vries EGE, van Vugt MATM, Fehrmann RSN. Transcriptional effects of copy number alterations in a large set of human cancers. Nat Commun 2020;11:715.

22. Subramanian A, Tamayo P, Mootha VK, Mukherjee S, Ebert BL, Gillette MA, et al. Gene set enrichment analysis: a knowledge-based approach for interpreting genome-wide expression profiles. Proc Natl Acad Sci 2005;102:15545–50.

23. Köhler S, Carmody L, Vasilevsky N, Jacobsen JOB, Danis D, Gourdine J-P, et al. Expansion of the human phenotype ontology (HPO) knowledge base and resources. Nucleic Acids Res 2019;47(D1):D1018–D1027.

24. Urzúa-Traslaviña CG, Leeuwenburgh VC, Bhattacharya A, Loipfinger S, van Vugt MATM, de Vries EGE, et al. Improving gene function predictions using independent transcriptional components. Nat Commun 2021;12:1464.

25. Canozo FJG, Zuo Z, Martin JF, Samee MdAH. Cell-type modeling in spatial transcriptomics data elucidates spatially variable colocalization and communication between cell-types in mouse brain. Cell Syst 2022;13:58-70.e5.

26. Bolton KL, Chen D, Fuente RIC de la, Fu Z, Murali R, Kbel M, et al. Molecular subclasses of clear cell ovarian carcinoma and their impact on disease behavior and outcomes. Clin Cancer Res 2022;28:4947–56.

27. Huang RY, Chen GB, Matsumura N, Lai HC, Mori S, Li J, et al. Histotype-specific copy-number alterations in ovarian cancer. BMC Méd Genom 2012;5:47.

28. Jongsma APM, Piek JMJ, Zweemer RP, Verheijen RHM, Gebbinck JWTK, Kamp GJ van, et al. Molecular evidence for putative tumour suppressor genes on chromosome 13q specific to BRCA1 related ovarian and fallopian tube cancer. Mol Pathol 2002;55:305.

29. Ko J, Fuccillo MV, Malenka RC, Südhof TC. LRRTM2 functions as a neurexin ligand in promoting excitatory synapse formation. Neuron 2009;64:791–8.

30. de Wit J, Sylwestrak E, O’Sullivan ML, Otto S, Tiglio K, Savas JN, et al. LRRTM2 interacts with neurexin1 and regulates excitatory synapse formation. Neuron 2009;64:799–806.

31. Ried T, Rudy B, Miera EV-S de, Lau D, Ward DC, Sen K. Localization of a highly conserved human potassium channel gene (NGK2-KV4; KCNC1) to chromosome 11p15. Genomics 1993;15:405–11.

32. Willis M, Trieb M, Leitner I, Wietzorrek G, Marksteiner J, Knaus H-G. Small-conductance calcium-activated potassium type 2 channels (SK2, KCa2.2) in human brain. Brain Struct Funct 2017;222:973–9.

33. Bourdeau ML, Laplante I, Laurent CE, Lacaille JC. KChIP1 modulation of Kv4.3-mediated A-type K+ currents and repetitive firing in hippocampal interneurons. Neuroscience 2011;176:173–87.

34. Ehmsen JT, Liu Y, Wang Y, Paladugu N, Johnson AE, Rothstein JD, et al. The astrocytic transporter SLC7A10 (Asc-1) mediates glycinergic inhibition of spinal cord motor neurons. Sci Rep 2016;6:35592.

35. Nath A, Cosgrove PA, Mirsafian H, Christie EL, Pflieger L, Copeland B, et al. Evolution of core archetypal phenotypes in progressive high grade serous ovarian cancer. Nat Commun 2021;12:3039.

36. Hanahan D, Weinberg RA. Hallmarks of cancer: the next generation. Cell 2011;144:646–74.

37. Reavis HD, Chen HI, Drapkin R. Tumor Innervation: Cancer Has Some Nerve. Trends Cancer 2020;6:1059–67.

38. Gysler SM, Drapkin R. Tumor innervation: peripheral nerves take control of the tumor microenvironment. J Clin Investig 2021;131:e147276.

39. Radin DP, Tsirka SE. Interactions between tumor cells, neurons, and microglia in the glioma microenvironment. Int J Mol Sci 2020;21:8476.

40. Venkatesh HS, Morishita W, Geraghty AC, Silverbush D, Gillespie SM, Arzt M, et al. Electrical and synaptic integration of glioma into neural circuits. Nature 2019;573:539–45.

41. Preter KD, Vandesompele J, Heimann P, Yigit N, Beckman S, Schramm A, et al. Human fetal neuroblast and neuroblastoma transcriptome analysis confirms neuroblast origin and highlights neuroblastoma candidate genes. Genome Biol 2006;7:R84.

42. Mark B, Lai S-L, Zarin AA, Manning L, Pollington HQ, Litwin-Kumar A, et al. A developmental framework linking neurogenesis and circuit formation in the Drosophila CNS. eLife 2021;10:e67510.

43. Allen JK, Armaiz-Pena GN, Nagaraja AS, Sadaoui NC, Ortiz T, Dood R, et al. Sustained adrenergic signaling promotes intratumoral innervation through BDNF induction. Cancer Res 2018;78:3233–3242.

44. Rains SL, Amaya CN, Bryan BA. Beta-adrenergic receptors are expressed across diverse cancers. Oncoscience 2017;4:95–105.

45. Eng JWL, Kokolus KM, Reed CB, Hylander BL, Ma WW, Repasky EA. A nervous tumor microenvironment: the impact of adrenergic stress on cancer cells, immunosuppression, and immunotherapeutic response. Cancer Immunol Immunother 2014;63:1115–28.

46. Drilon A, Siena S, Ou S-HI, Patel M, Ahn MJ, Lee J, et al. Safety and antitumor activity of the multitargeted Pan-TRK, ROS1, and ALK inhibitor entrectinib: combined results from two phase I trials (ALKA-372-001 and STARTRK-1). Cancer Discov 2017;7:400–9.

47. Ardini E, Menichincheri M, Banfi P, Bosotti R, Ponti CD, Pulci R, et al. Entrectinib, a pan–TRK, ROS1, and ALK inhibitor with activity in multiple molecularly defined cancer indications. Mol Cancer Ther 2016;15:628–39.

48. Drilon A, Laetsch TW, Kummar S, DuBois SG, Lassen UN, Demetri GD, et al. Efficacy of larotrectinib in TRK fusion–positive cancers in adults and children. N Engl J Med 2018;378:731–9.

49. Burris HA, Shaw AT, Bauer TM, Farago AF, Doebele RC, Smith S, et al. Abstract 4529: Pharmacokinetics (PK) of LOXO-101 during the first-in-human phase I study in patients with advanced solid tumors: Interim update. Cancer Res 2015;75:4529–4529.

50. Barr JL, Kruse A, Restaino AC, Tulina N, Stuckelberger S, Vermeer SJ, et al. Intra-tumoral nerve-tracing in a novel syngeneic model of high-grade serous ovarian carcinoma. Cells 2021;10:3491.

51. Restaino AC, Walz A, Vermeer SJ, Barr J, Kovács A, Fettig RR, et al. Functional neuronal circuits promote disease progression in cancer. Sci Adv 2023;9:eade4443.

52. Balood M, Ahmadi M, Eichwald T, Ahmadi A, Majdoubi A, Roversi K, et al. Nociceptor neurons affect cancer immunosurveillance. Nature. 2022;611:405–12.

53. Li X, Peng X, Yang S, Wei S, Fan Q, Liu J, et al. Targeting tumor innervation: premises, promises, and challenges. Cell Death Discov 2022;8:131.

54. Padmanaban V, Keller I, Seltzer ES, Ostendorf BN, Kerner Z, Tavazoie SF. Neuronal substance-P drives breast cancer growth and metastasis via an extracellular RNA-TLR7 axis. bioRxiv 2024;2024.03.08.584128.

55. Williams K. Ifenprodil, a novel NMDA receptor antagonist⍰: site and mechanism of action. Curr Drug Targets 2001;2:285–98.

56. Reid JG, Gitlin MJ, Altshuler LL. Lamotrigine in psychiatric disorders. J Clin Psychiatry 2013;74:675–84.

57. Ramamoorthy MD, Kumar A, Ayyavu M, Dhiraviam KN. Reserpine induces apoptosis and cell cycle arrest in hormone independent prostate cancer cells through mitochondrial membrane potential failure. Anti-Cancer Agents Med Chem 2019;18:1313–22.

58. North WG, Liu F, Tian R, Abbasi H, Akerman B. NMDA receptors are expressed in human ovarian cancer tissues and human ovarian cancer cell lines. Clin Pharmacol⍰: Adv Appl 2015;7:111–7.

