## Supplementary figures tables and methods for "Transcriptional pattern enriched for synaptic signaling is associated with shorter survival of patients with high-grade serous ovarian cancer": Supplementary methods 24022025.docx

**Data acquisition**

Publicly available raw microarray bulk transcriptomics were extracted from the Gene Expression Omnibus (GEO)[1]. We confined to the Affymetrix HG-U133 Plus 2.0 platform (GEO accession identifier: GPL570). Samples were selected for analysis if they represented epithelial ovarian cancer (EOC) obtained from patients. Also, samples were collected from non-cancerous ovarian tissue obtained with surface brushings or laser micro-dissected ovarian epithelium (n = 25), ovarian epithelial cells isolated using short-term culture (n = 9) or derived from complete ovaries (n = 9). Cell line samples were excluded. For all selected samples, relevant clinicopathological data were collected whenever available, including grade, stage, age, treatment history, debulking status, progression-free survival (PFS), and OS.

*CCLE dataset*

Raw bulk transcriptomic data was obtained from the CCLE project, which conducted a detailed genetic characterization of a large panel of human cancer cell lines[2]. Expression data within the CCLE project was generated with Affymetrix HG-U133 Plus 2.0. This dataset is referred to as the CCLE dataset throughout this manuscript.

*GDSC dataset*

From the GDSC portal, we obtained raw expression data generated with Affymetrix HG-U219[3]. The aim of the GDSC project is to identify molecular features of cancer that predict response to anti-cancer drugs. This dataset is referred to as the GDSC dataset throughout this manuscript.

TCGA dataset

From TCGA, we obtained the pre-processed and normalized level 3 RNA-seq (version 2) data for 27 cancer datasets available at the Broad GDAC Firehose portal (downloaded January 2017 https://gdac.broadinstitute.org/). For each sample, we downloaded RNA-Seq with Expectation Maximization (RSEM) gene normalized data (identifier: illuminahiseq_rnaseqv2 RSEM_genes_normalized)[4].

Spatial transcriptomic profiles

From 10xGenomics repository, we sourced the spots-without-tissue-filtered Spatial transcriptomic profiles of three samples from ovarian cancer patients in h5 format. Stained images of these three samples along with scale factors and tissue positions were also downloaded from the same repository. These three samples were collected from patients had high grade serous ovarian cancer, serous papillary carcinoma and endometroid adenocarcinoma of the ovary respectively. The repositories are mentioned in the following:

1. <https://www.10xgenomics.com/datasets/human-ovarian-cancer-11-mm-capture-area-ffpe-2-standard>
2. <https://www.10xgenomics.com/datasets/human-ovarian-cancer-1-standard>
3. <https://www.10xgenomics.com/datasets/human-ovarian-cancer-whole-transcriptome-analysis-stains-dapi-anti-pan-ck-anti-cd-45-1-standard-1-2-0>

We also collected publicly available spatial resolved transcriptomic profiles from eight ovarian cancer samples sourced from GEO (study ID GSE211956).

**Sample processing and quality control**

Non-corrupted raw data CEL files were downloaded from GEO for the selected samples. To identify samples that were uploaded to GEO multiple times, we generated an MD5 hash for each CEL file. After the removal of duplicate CEL files, preprocessing and aggregation of CEL files was performed with robust multiarray averaging (RMA using the justRMA function from R package aroma.affymetrix v3.2.0) method using R version 3.5.2. Quality control was performed using principal component analysis as previously described[5]. MD5 hash duplicate removal does not detect identical samples when meta-data is different. Pearson correlation coefficients among bulk transcriptomics were obtained to identify samples with identical expression values. One sample from each set of duplicate samples was randomly chosen, and the rest were removed from subsequent analyses.

**Consensus independent component analysis (c-ICA)**

The bulk transcriptomics included in this analysis were generated with complex tumour tissues. These tissues contain a complex mixture of heterogeneous tumour cells and non-tumour cells present in the tumour microenvironment. Therefore, the resulting profiles represent the average transcriptomic patterns of cells present in the biopsies. c-ICA was utilized to segregate the average transcriptomic patterns of complex biopsies into statistically independent transcriptional components[6].

Applying ICA on a bulk transcriptomic dataset with 𝑝 genes and 𝑛 samples results in the extraction of *i* independent components of dimension 1×*p* (hereafter called estimated sources, ESs) and a mixing matrix (MM) of dimension *i* × *n* which contains the coefficients of ESs in each sample. The weight of each ES represents the direction and magnitude of its effect on the expression level of each gene, and the coefficients of MM represent activity scores of the ESs in the corresponding sample. In ICA, a preprocessing technique called whitening is applied to the input dataset to make the estimation more time efficient. Whitening was used to transform bulk transcriptomics of all samples so that the transformed profiles are uncorrelated and have a variance of one. Next, ICA was performed on the whitened dataset using the FastICA function from the FastICA package (version 1.2.0), resulting in the extraction of *i* independent components and a mixing matrix. The parameter *i* was chosen as the number of top principal components, which captured 90% of the total variance seen in the whitened dataset.

In ICA, an initial random weight vector with a variance of 1 has to be chosen to obtain statistically independent ESs. Hence, different initial random weight vectors could result in different sets of ESs. To retrieve a set of consensus ESs (hereafter called transcriptional components, TCs), we performed 25 ICA runs, each with a different random initialization weight vector. The assumption is that over a large number of runs of ICA, fastICA algorithm does not converge to any local solution for most of the runs. ESs extracted from these runs were clustered together if the absolute value of the Pearson correlation between them was > 0.9. Clusters with ESs from >50% of the runs were used to obtain *m* TCs using the following formula:

$${TC}_{p\times1}=(1/n)\sum_{i=1}^{n} ({ES}_{i}\times sign\left( correlation\left( {ES}_{1},{ES}_{i} \right) \right))$$

These *m* TCs and the bulk transcriptomic dataset with *p* genes and *n* samples ($X_{p\times n}$) was used in the below formula to obtain consensus mixing matrix (or CMM) which contains the coefficients of TCs in each sample:

$${CMM}_{m\times n}=(\left( {TC}^{'} \right)_{m\times p}\times{TC}_{p\times m})^{-1}\times{({TC}^{'})}_{m\times p}\times X_{p\times n}$$

Similar to the MM, the coefficients of CMM represent activity scores of the TCs in the corresponding sample.

**Univariate survival analysis**

As the coefficients of CMM for each TC vary from one sample to another, we hypothesized that these activity scores of the TCs might be associated with OS. To assess this association, we calculated the -log_10_(p-value) from Cox regression for each TC_i_ as predictor, denoted as 'original_minus_log10_p_i_'. To reduce the probability of false-positive association due to multiple testing, we conducted a permutation test. First, the sample labels of the TC activities in the MM were permutated 10,000 times, to ensure that the correlation structure remained intact, resulting in 10,000 sets of permutated TCs. Then, on each TC_i_ separately the following steps were conducted:

1. Cox regression is performed using the permutated TC_i_ activities.
2. From the analyses of step 1, -log10(p-value) corresponding to all permuted TCs are obtained (permuted_minus_log10_p).
3. Sort the values of permuted_minus_log10_p in decreasing order (sp_minus_log10_p).
4. Sort the values of original_minus_log10_p in decreasing order (so_minus_log10_p).
5. For every *j*-th value of so_minus_log10_p (so_minus_log10_p_j_), obtain the number of values of sp_minus_log10_p_i_ greater than so_minus_log10_p_j_, defined as *f_j_*.
6. We subset so_minus_log10_p till the j-th entry where *f_j_* / j > 1% (false discovery rate of 1%).
7. Obtain the optimal cutoff for defining false discoveries of sp_minus_log10_p_i_ (*oc_i_*) as the maximum value of so_minus_log10_p.

Next, we defined a single optimal cutoff of *oc* from all the TCs, as 80% quantile of values of occ (confidence level of 80%). For each TC_i_, if original_minus_log10_p_i_ >= the single optimal cutoff, it was considered statistically significantly associated with OS in the permutation test framework.

**Multivariate survival analysis**

Next, we performed multivariate Cox proportional hazards analysis using the known prognostic parameters age, stage, debulking status, and grade along with the TCs one at a time as predictors. A smaller subset of samples containing information about all of these clinicopathological variables was used for this analysis. For each TC_i_, multivariate survival analysis with OS as the outcome variable was conducted using the following steps:

1. -log_10_(adjusted p-value) for each TC_i_ as a predictor (original_minus_log10_adj_p_i_) were obtained from the multivariate survival analysis.
2. Permutation tests were performed on each TC_i_ exactly as described in the univariate survival analysis section. As a result, we obtained a list of TCs which are statistically significantly associated with OS after removing the effects of age, stage, debulking status, and grade in the permutation test framework.

**Survival tree analysis**

Survival tree analysis is used to classify samples with clinicopathological information based on the difference between survival probabilities at certain time intervals. A subset of platinum-treated SOC samples with follow-up information was used in survival tree analysis. All TCs associated with OS in the univariate or multivariate survival analyses served as classifiers along with all clinicopathologic parameters (age, stage, debulking status, and grade). Survival tree analysis was performed using the following steps:

1. For each of the classifiers (C_i_):
   a. If the classifier is a numeric variable,
      i. For each possible weight of the classifier as a cutoff ( c ):
         1. Obtain two subsets of the samples.
            a. Samples having weights < cutoff weight
            b. Or samples having weights ≥ cutoff weight
         2. Obtain survival probabilities for both of the subsets.
         3. Compare the survival probabilities using the log-rank test and obtain the log-rank statistic (LRC_i,c_).
      ii. Obtain the optimum cutoff c (oc_i_) for which log-rank statistic LRC_i,c_ is maximum. Assign maximum of LRC_i,c_ as max_LRC_i_.
   b. If the classifier is a categorical variable,
      i. For each possible combination of different levels of the classifier:
         1. Obtain two subsets of samples A and B.
         2. Obtain survival probabilities for both of the subsets.
         3. Compare the survival probabilities using the log-rank test and obtain the log-rank statistic (LRC_i,AB_).
      ii. Obtain the optimum combination of levels in two subsets A, B (oc_i_) for which log-rank statistic LRC_i,AB_ is maximum. Assign maximum of LRC_i,AB_ as max_LRC_i_.
2. Obtain the most significant classifier along with the optimum cutoff/combination of levels (oc) for which max_LRC_i_ is maximum among all classifiers.
3. Classify the samples into two subsets (subset_1 and subset_2) using the most significant classifier and optimum cutoff/combination of levels.
4. In each of the subsets of the dataset, repeat steps 1, 2 & 3 till the following constraints are maintained.
   a. Number of samples in subset_1 + number of samples in subset_2 ≥ 50
   b. number of uncensored events in subset_1 + number of uncensored events in subset_2 ≥ 25
   c. number of samples in subset_1 or number of samples in subset_2 ≥ 17

After that, 10,000 iterations of the above four steps were performed using random 80% of the samples each time to investigate the robustness of the significance of the classifiers obtained from the survival tree analysis mentioned above.

To assess the goodness-of-fit of the survival tree, the following steps were conducted:

1. A new variable called survival cohorts was created to store the terminal node number in the survival tree of each sample.
2. Univariate Cox regression with this new variable survival cohorts as a predictor was performed to assess its association with OS.
3. Concordance statistic for this Cox regression model was obtained to evaluate the classification power of the survival tree.

To quantify the robustness of the survival tree, the following steps were conducted:

1. Survival tree analysis was conducted 20,000 times using random 80% of the samples each time.
2. For the survival tree with all samples
   a. For each classifier *C*
      i. NOS_node*_C_*_,_*_n_* was obtained as the number of samples present in node *n* where *C* was the most significant classifier.
      ii. Max_NOS_node*_C_* was obtained as maximum of NOS_node*_C_*_,_*_n_*.
      iii. R_all_samples_survival_tree*_C_* was obtained as rank of Max_NOS_node*_C_*.
3. For each of the 20,000 iterations *i*
   a. For each classifier *C*
      i. NOS_node*_C_*_,_*_n_* was obtained as the number of samples present in node *n* where *C* was the most significant classifier.
      ii. Max_NOS_node*_C_* was obtained as maximum of NOS_node*_C_*_,_*_n_*.
      iii. R*_C_*_,_*_i_* was obtained as the rank of Max_NOS_node*_C_*.
   b. Robustness_statistic*_i_* was obtained as the Spearman correlation coefficient between R*_C_*_,_*_I_* and R_all_samples_survival_tree*_C_*

The robustness of the survival tree was presented by the interquartile range and median of Robustness_statistic*_i_*_._

**Associating the identified transcriptional components with biological processes**

We characterized biological activity represented by a TC using multiple methods. Firstly, based on our recently published method called Transcriptional Adaptation to Copy Number Alterations (TACNA) profiling, we identified a subset of TCs that capture the downstream effects of CNAs on mRNA expression levels[5]. Secondly, we performed GSEA on each TC using gene set collections (n = 16) collected from The Human Phenotype Ontology (The Monarch Initiative), the Mammalian Phenotypes (Mouse Genome Database), and the Molecular Signatures Database (MsigDB)[7–9]. We included all gene sets with 10-500 genes after filtering out genes that were not present in the expression profiles. Enrichment of each gene set was tested according to the two-sample Welch’s t-test for unequal variance between the set of genes, which were under investigation, versus the set of genes that was not under investigation. To allow comparison between gene sets of different sizes, we transformed the Welch’s t statistic to a Z-score.

Finally, we used the genetICA-network (available at <http://www.genetica-network.com>), to predict the likely function of the top genes inside a component[10]. In short, a GBA approach was used to predict likely functions for genes based on gene co-regulation. For this, we conducted a consensus-ICA on an unprecedented scale. In short, a covariance matrix was calculated between 19,635 genes using the expression patterns of 106,462 bulk transcriptomics generated with Affymetrix HG-U133 Plus 2.0 representing the many disease states, cellular states, and genetic and chemical perturbations that were obtained. Consensus-ICA was performed on the covariance matrix. This identified a large set of CESs and a mixing matrix reflecting the activity of each source in the expression pattern of the gene across the samples. Next, a GBA approach was used to predict the functionality of individual genes. First, we retrieved 16 public gene set collections describing a large range of biological processes and phenotypes. For each gene set, we calculated its ‘bar code’ by averaging the MM weight of its member genes. Next, for each gene in the MM, the distance correlation was determined between its MM weights and the gene set bar code. A high correlation between a gene’s MM weight and a gene set bar code indicated that the gene under investigation shared functionality with the genes of the specific gene set under investigation. Significance levels were obtained with permutated data (250 permutations). This strategy was used on 23,372 well-described functional gene sets, which enabled us to create a comprehensive network of predicted functionalities of individual genes. From each studied TC, we separately selected the top and bottom 250 genes with a weight of ≥ 3 or ≤ -3 and created co-functionality networks using a network threshold of 0.65. For all resulting gene clusters consisting of ≥ 5 genes, the top 10 gene sets with a mean Z-score >2 were used for interpretation of the biological signal within a TC.

**Cross-study transcriptional component projection**

A cross-study transcriptional component projection was conducted to investigate the activity scores of the transcriptional components in independently obtained mRNA expression profiles. Following steps were conducted for each cross-study transcriptional component projection analysis where transcriptional components (TC) of dataset $i$ were used to obtain activity scores of the same TC’s in samples of dataset $j$:

1. Genes not present in both dataset $i$ and $j$ were removed from the analysis.
2. Both dataset $i$ and dataset $j$ were standardized on gene-level separately, which means each gene expression is transformed to a mean of zero and standard deviation of one.
3. Both of these standardized datasets were sample-wise merged to obtain a combined dataset ($Combined\_i\_used\_for\_j$).
4. Transcriptional component matrix of dataset $i$ ($\mathrm{TC}_{i}$) and $Combined\_i\_used\_for\_j$ were used to obtain consensus mixing matrix ($CMMcombined\_i\_used\_for\_j$). $CMMcombined\_i\_used\_for\_j=((\mathrm{TC}_{i})'\times\mathrm{TC}_{i})^{-1}\times(\mathrm{TC}_{i})'\times Combined\_i\_used\_for\_j$
5. The subset of coefficients $CMMcombined\_i\_used\_for\_j$ for samples from dataset j is obtained as $CMM\_j$

**Identification of significant activity of each TC in spatial transcriptomic profiles**

To identify if the activity scores of each TC in each spatial transcriptomic profile is significantly different from the null distribution of possible activity scores, we conducted the following steps:

- A set of 3,000 permutations of all the genes weights in the TC i were conducted (ith permuted TC is denoted by $TC\_permuted\_i$). For each permutation r:
  a. Obtain the permuted activity score for the sample s using the following formula:

$${CMM\_permuted}_{r}=(\left( {TC\_permuted\_i}^{'} \right)_{1\times p}\times{TC\_permuted\_i}_{p\times1})^{-1}\times{({TC\_permuted\_i}^{'})}_{1\times p}\times X_{p\times n}$$

b. Thereafter a Johnson transformation was conducted on the vector $CMM\_permuted$ so that the distribution was as similar as possible to normal distribution with mean zero and standard deviation of one (referred to as $CMM\_permuted\_Johnsontransformed$). The same transformation was applied on the original CMMi,s.
c. Finally a p-value was obtained based on the position of the Johnson transformed CMMi,s with respect to the generalized normal distribution fitted to the $CMM\_permuted\_Johnsontransformed$
d. Log-transformed p-values were thereafter plotted in a heatmap incorporating the row and column position of the individual spatial transcriptomic profiles to visualize the location of significant activity of the TCs.

**Determination of spatial transcriptomic profiles’ significant activity locations for individual transcriptional component**

Public spatial resolved transcriptomic profiles from three ovarian cancer samples, generated using the 10xGenomics Visium platform, were obtained for analysis. The samples were from patients with high grade serous, serous papillary and endometrioid ovarian cancer respectively. Activity for each TC across every location within the spatial samples was ascertained through the cross-study projection methodology referred to in the previous method section. To discern the markedly active areas within the spatial samples for each TC, we incorporated a permutation-driven approach. We derived a null-distribution of activities for each TC-location pairing by performing 3,000 permutations and subsequent projection. The p-value of the observed activity was set as the number of permutations that gave a higher activity divided by the number of permutations. This p-value quantifies the significance of the deviation of the TC's activity at a given location from its baseline null distribution. After this, we visualized the p-values, post their log transformation, using a heatmap. This visualization aided in highlighting the areas with notable activity, aligned against the stained representation of the tissue sample.

**Colocalization analysis of TC activities in spatial transcriptomics**

The colocalization method is based on the principles of colocalization analysis in microscopy images and is adapted from a previously published colocalization analysis.[11] We first chose highly (in)active regions by selecting spots where the TCs activity of the permutation-corrected mixing matrix of the spatial transcriptomic images was above 2 for highly active or below -2 for inactive TC processes. Then, for each image and TC, kernel density estimation was performed to estimate the density of highly active or inactive regions using the R package ks 1.14.1.[12] Only TCs with at least 20 active or inactive spots were considered. The optimal kernel bandwidth was determined using the least-squares cross-validation bandwidth selector with the method "Hlscv.diag" and an initial bandwidth matrix of [(9,0),(0,9)]. The binned background grid (bgridsize) was defined as the dimensions of the spots in the spatial transcriptomic image. The kernel was then fitted using the "kde" function with weights being the absolute TC activity per spot of the (in)active regions. Next, we retained only those locations where the kernel density regions exceeded the 75^th^ percentile of the kernel densities. We then assessed the similarity of kernel densities for all combinations of active and inactive regions of TCs of an image by taking the union of the selected regions and calculating the Pearson correlation coefficients *ρ* between the density values of two TCs. A *ρ* of 1 indicates that the (in)active regions of two TCs are colocalized, while a *ρ* of -1 indicates that they are spatially exclusive. To enhance the colocalization analysis between TCs, we considered both coexisting and repelling properties of (in)active processes. To quantify this, we implemented a colocalization score between two TCs *a* and *b* and their respective active and inactive regions as follows:

colocalization score_a,b_ = ( *ρ_a_active, b_active_* - *ρ_a_active, b_*__inactive_ ) - ( *ρ_a_inactive, b_active_* - *ρ_a_inactive, b_*__inactive_ )

A colocalization score of 4 between two TCs indicates that the active and inactive regions of these TCs are colocalized, with the inactive regions of one TC being spatially exclusive to the active regions of the other TC and vice versa. Conversely, a colocalization score of -4 between two TCs also indicates colocalization, but with reversed activity, i.e. the active regions of one TC are colocalized with the inactive regions of the other TC or vice versa. A colocalization score close to 0 implies that the activities of two TCs are spatially exclusive.

**Cross-study projection of TCs on single cell transcriptional profiles**

We analyzed single-cell transcriptional data consisting of 71,965 cell-type-annotated profiles from the study on patients with HGSOC (study id GSE158722). Of these, 63,793 profiles were categorized into 21 distinct cell types. To explore the remaining unannotated profiles, we randomly selected 10% for further analysis. We first removed the first principal component from the profile correlation matrix to minimize pervasive background signals. We then conducted a cross-dataset projection of TCs onto these profiles, calculating activity scores across diverse cellular contexts.

**Univariate survival analysis using transcriptional profiles of patients with ovarian clear cell carcinoma**

Expression profiles and survival data for ovarian clear cell carcinoma (OCCC) patients from Bolton et al. were obtained from the study's GitHub repository: <https://github.com/kbolton-lab/Bolton_OCCC>.[13]  Next, we calculated the activity scores for all TCs across all samples after performing preprocessing as described in above. Subsequently, we conducted the univariate association analysis, as outlined earlier, using survival information from 120 patients for whom transcriptional profiles were available. The results are summarized by displaying log-transformed p-values along with the signs of the coefficients.

**References**

1. Clough, E.; Barrett, T. Statistical Genomics, Methods and Protocols. *Methods Mol. Biol.* **2016**, *1418*, 93–110, doi:10.1007/978-1-4939-3578-9_5.

2. Barretina, J.; Caponigro, G.; Stransky, N.; Venkatesan, K.; Margolin, A.A.; Kim, S.; Wilson, C.J.; Lehár, J.; Kryukov, G.V.; Sonkin, D.; et al. The Cancer Cell Line Encyclopedia Enables Predictive Modelling of Anticancer Drug Sensitivity. *Nature* **2012**, *483*, 603–607, doi:10.1038/nature11003.

3. Yang, W.; Soares, J.; Greninger, P.; Edelman, E.J.; Lightfoot, H.; Forbes, S.; Bindal, N.; Beare, D.; Smith, J.A.; Thompson, I.R.; et al. Genomics of Drug Sensitivity in Cancer (GDSC): A Resource for Therapeutic Biomarker Discovery in Cancer Cells. *Nucleic Acids Res.* **2013**, *41*, D955–D961, doi:10.1093/nar/gks1111.

4. *GDC Data User’s Guide NCI Genomic Data Commons (GDC)*;

5. Bhattacharya, A.; Bense, R.D.; Urzúa-Traslaviña, C.G.; Vries, E.G.E. de; Vugt, M.A.T.M. van; Fehrmann, R.S.N. Transcriptional Effects of Copy Number Alterations in a Large Set of Human Cancers. *Nat. Commun.* **2020**, *11*, 715, doi:10.1038/s41467-020-14605-5.

6. Chiappetta, P.; Roubaud, M.C.; Torrsani, B. Blind Source Separation and the Analysis of Microarray Data. *J. Comput. Biol.* **2004**, *11*, 1090–1109, doi:10.1089/cmb.2004.11.1090.

7. Liberzon, A.; Subramanian, A.; Pinchback, R.; Thorvaldsdóttir, H.; Tamayo, P.; Mesirov, J.P. Molecular Signatures Database (MSigDB) 3.0. *Bioinformatics* **2011**, *27*, 1739–1740, doi:10.1093/bioinformatics/btr260.

8. Liberzon, A.; Birger, C.; Thorvaldsdóttir, H.; Ghandi, M.; Mesirov, J.P.; Tamayo, P. The Molecular Signatures Database Hallmark Gene Set Collection. *Cell Syst.* **2015**, *1*, 417–425, doi:10.1016/j.cels.2015.12.004.

9. Subramanian, A.; Tamayo, P.; Mootha, V.K.; Mukherjee, S.; Ebert, B.L.; Gillette, M.A.; Paulovich, A.; Pomeroy, S.L.; Golub, T.R.; Lander, E.S.; et al. Gene Set Enrichment Analysis: A Knowledge-Based Approach for Interpreting Genome-Wide Expression Profiles. *Proc. Natl. Acad. Sci.* **2005**, *102*, 15545–15550, doi:10.1073/pnas.0506580102.

10. Urzúa-Traslaviña, C.G.; Leeuwenburgh, V.C.; Bhattacharya, A.; Loipfinger, S.; Vugt, M.A.T.M. van; Vries, E.G.E. de; Fehrmann, R.S.N. Improving Gene Function Predictions Using Independent Transcriptional Components. *Nat. Commun.* **2021**, *12*, 1464, doi:10.1038/s41467-021-21671-w.

11. Canozo, F.J.G.; Zuo, Z.; Martin, J.F.; Samee, Md.A.H. Cell-Type Modeling in Spatial Transcriptomics Data Elucidates Spatially Variable Colocalization and Communication between Cell-Types in Mouse Brain. *Cell Syst.* **2022**, *13*, 58-70.e5, doi:10.1016/j.cels.2021.09.004.

12. Chacón, J.E.; Duong, T. Multivariate Kernel Smoothing and Its Applications. **2018**, 89–110, doi:10.1201/9780429485572-14.

13. Bolton, K.L.; Chen, D.; Fuente, R.I.C. de la; Fu, Z.; Murali, R.; K�bel, M.; Tazi, Y.; Cunningham, J.M.; Chan, I.C.C.; Wiley, B.J.; et al. Molecular Subclasses of Clear Cell Ovarian Carcinoma and Their Impact on Disease Behavior and Outcomes. *Clin. Cancer Res.* **2022**, *28*, 4947–4956, doi:10.1158/1078-0432.ccr-21-3817.
