## Supplementary figures tables and methods for "Transcriptional pattern enriched for synaptic signaling is associated with shorter survival of patients with high-grade serous ovarian cancer": Supplementary Table S1.docx

**Supplementary Table S1.** Patient characteristics. Abbreviations: NA = not applicable, high-grade SOC = high-grade serous ovarian cancer, LGSOC = low-grade serous ovarian cancer, LMP = low malignant potential, OCCC = ovarian clear cell cancer.

|  | All samples  n = 1125 | % | Univariate  n = 541 | % | Multivariate & survival tree  n = 301 | % |
| --- | --- | --- | --- | --- | --- | --- |
| Age |  |  |  |  |  |  |
| mean (range) | 58.0 (22-89) | *52* | 59.6 (22-89) | *75* | 59.7 (23-89) | *100* |
| NA | 539 | *48* | 133 | *25* |  |  |
| Grade |  |  |  |  |  |  |
| 1 | 45 | *4.0* | 27 | *5.0* | 7 | *2.3* |
| 2 | 161 | *14* | 141 | *26* | 92 | *31* |
| 3 | 460 | *41* | 336 | *62* | 202 | *67* |
| 4 | 19 | *1.7* | 15 | *2.8* |  |  |
| NA | 440 | *39* | 22 | *4.1* |  |  |
| Stage |  |  |  |  |  |  |
| 1 | 147 | *13* | 45 | *8.3* | 18 | *6.0* |
| 2 | 76 | *6.8* | 30 | *5.5* | 14 | *4.7* |
| 3 | 538 | *48* | 358 | *66* | 235 | *78* |
| 4 | 93 | *8.3* | 55 | *10* | 34 | *11* |
| NA | 271 | *24* | 53 | *9.8* |  |  |
| Subtype |  |  |  |  |  |  |
| HGSOC | 678 | *60* | 470 | *87* | 294 | *97* |
| LGSOC | 40 | *3.6* | 11 | *2.0* | 7 | *2.3* |
| LMP | 53 | *4.7* | 19 | *3.5* |  |  |
| Serous - undefined | 70 | *6.2* | 2 | *0.4* |  |  |
| Endometrioid | 110 | *9.8* | 25 | *4.6* |  |  |
| OCCC | 96 | *8.5* | 6 | *1.1* |  |  |
| Mucinous | 35 | *3.1* | 8 | *1.5* |  |  |
| Normal | 43 | *3.8* |  |  |  |  |
| Platinum-treated |  |  |  |  |  |  |
| Yes | 523 | *47* | 440 | *81* | 301 | *100* |
| No | 52 | *4.6* | 49 | *9.1* |  |  |
| Unknown | 550 | *49* | 52 | *9.6* |  |  |
| Taxol-treated |  |  |  |  |  |  |
| Yes | 406 | *36* | 323 | *60* | 235 | *78* |
| No | 169 | *15* | 160 | *30* | 66 | *22* |
| Unknown | 550 | *49* | 52 | *10* |  |  |
| Debulking status |  |  |  |  |  |  |
| Optimal | 303 | *27* | 303 | *56* | 162 | *54* |
| Suboptimal | 187 | *17* | 184 | *34* | 139 | *46* |
| NA | 635 | *56* | 54 | *10* |  |  |
