## Supplementary figures tables and methods for "Transcriptional pattern enriched for synaptic signaling is associated with shorter survival of patients with high-grade serous ovarian cancer": Supplementary Table S2.docx

**Supplementary Table S2.** Overview of the number of samples from each GEO series and the corresponding study. Abbreviations: PMID = PubMed identification number.

| Series | # samples | PMID | Reference |
| --- | --- | --- | --- |
| GSE105437 | 15 | 29212026 | Noh et al. 2017 |
| GSE107931 | 4 | 29339543 | Curry et al. 2018 |
| GSE10971 | 11 | 18593983 | Tone et al. 2008 |
| GSE115635 | 4 | 29860390 | Yeung et al. 2019 |
| GSE12172 | 11 | 19010816 | Anglesio et al. 2008 |
| GSE14001 | 23 | 19525924 | Tung et al. 2009 |
| GSE14407 | 20 | 20040092 | Bowen et al. 2009 |
| GSE15578 | 6 | 19956396 | Pejovic et al. 2009 |
| GSE18521 | 59 | 19962670 | Mok et al. 2009 |
| GSE19352 | 17 | 20179205 | Iorio et al. 2010 |
| GSE19829 | 28 | 20547991 | Konstantinopoulos et al. 2010 |
| GSE20565 | 90 | 20492709 | Meyniel et al. 2010 |
| GSE2109 | 137 | - | *The International Genomics Consortium. 2019*; Available from:  <https://www.ncbi.nlm.nih.gov/bioproject/PRJNA91763>. |
| GSE26193 | 14 | 22101765 | Mateescu et al. 2011 |
| GSE27651 | 25 | 21451362 | King et al. 2011 |
| GSE27659 | 10 | 20802181 | Wong et al. 2010 |
| GSE29450 | 10 | 21754983 | Stany et al. 2011 |
| GSE32062 | 10 | 22241791 | Yoshihara et al. 2012 |
| GSE3526 | 4 | 16572319 | Roth et al. 2006 |
| GSE36668 | 4 | 23029477 | Elgaaen et al. 2012 |
| GSE39204 | 60 | 23340297 | Kaoru Abiko et al. 2013 |
| GSE40595 | 3 | 23824740 | Yeung et al. 2013 |
| GSE44104 | 60 | 23934190 | Wu et al. 2014 |
| GSE51373 | 28 | 24237932 | Koti et al. 2013 |
| GSE52460 | 3 | 24666724 | Hill et al. 2014 |
| GSE54388 | 5 | 28199976 | Yeung et al. 2017 |
| GSE55512 | 12 | 25867264 | K Abiko et al. 2015 |
| GSE63885 | 94 | 24478986 | Lisowska et al. 2014 |
| GSE65986 | 55 | 26147301 | Uehara et al. 2015 |
| GSE69428 | 10 | 26415052 | Yamamoto et al. 2016 |
| GSE73168 | 8 | 30710055 | Gao et al. 2019 |
| GSE9899 | 285 | 18698038 | Tothill et al. 2008 |
