## Supplementary figures tables and methods for "Transcriptional pattern enriched for synaptic signaling is associated with shorter survival of patients with high-grade serous ovarian cancer": Supplementary Table S5.docx

**Supplementary Table S5.** Clinicopathological parameters per survival tree node in platinum-treated high-grade serous ovarian cancer.

| Node 1 (TC 121 / Neuronal development) |  | Low (#) | % | High (#) | % |
| --- | --- | --- | --- | --- | --- |
| Stage | 1  2  3  4 | 14  12  203  30 | *5.4*  *4.6*  *78*  *12* | 2  1  29  3 | *5.7*  *2.9*  *83*  *8.6* |
| Age, mean (range) |  | 59 (23-89) | | 64 (33-79) | |
| Taxol treated | Yes  No | 207  52 | *80*  *20* | 23  12 | *66*  *34* |
| Neo-adjuvant | Yes  No | 11  248 | *4.2*  *96* | 1  34 | *2.9*  *97* |
| Debulking | Optimal  Suboptimal | 140  119 | *54*  *46* | 18  17 | *51*  *49* |
| Series | 19829  20565  26193  9899 | 26  54  10  169 | *10*  *21*  *3.9*  *65* | 1  5  1  28 | *2.9*  *14*  *2.9*  *80* |
| 1 year survival (95% CI) | 95% (93-98) | | | 86% (75-98) | |
| 3 year survival (95% CI) | 62% (56-69) | | | 32% (19-54) | |
| 5 year survival (95% CI) | 38% (31-46) | | | 14% (5-43) | |
| Median survival | 1391 (1251-1647) | | | 671 (610-1159) | |

A. Clinicopathological parameters for subgroups defined by node 1 (TC 121 / Neuronal development)

| Node 2 (Stage) |  | 1-2-3 (#) | % | 4 (#) | % |
| --- | --- | --- | --- | --- | --- |
| Stage | 1  2  3  4 | 14  12  203 | *6.1*  *5.2*  *89* | 30 | *100* |
| Age, mean (range) |  | 60 (23-89) | | 58 (38-80) | |
| Taxol treated | Yes  No | 185  44 | *81*  *19* | 22  8 | *73*  *27* |
| Neo-adjuvant | Yes  No | 6  223 | *2.6*  *97* | 5  25 | *17*  *83* |
| Debulking | Optimal  Suboptimal | 132  97 | *58*  *42* | 8  22 | *27*  *73* |
| Series | 19829  20565  26193  9899 | 22  46  8  153 | *9.6*  *20*  *3.5*  *67* | 4  8  2  16 | *13*  *27*  *7*  *53* |
| 1 year survival (95% CI) | 96% (94-99) | | | 93% (84-100) | |
| 3 year survival (95% CI) | 67% (60-74) | | | 27% (14-53) | |
| 5 year survival (95% CI) | 40% (33-49) | | | 0 (NA-NA) | |
| Median survival (95% CI) | 1454 (1342-1769) | | | 903 (732-1464) | |

B. Clinicopathological parameters for subgroups defined by node 2 (Stage)

| Node 3 (Stage) |  | 1-2 (#) | % | 3 (#) | % |
| --- | --- | --- | --- | --- | --- |
| Stage | 1  2  3 | 14  12 | *54*  *46* | 203 | *100* |
| Age, mean (range) |  | 59 (43-77) | | 60 (23-89) | |
| Taxol treated | Yes  No | 19  7 | *73*  *27* | 166  37 | *82*  *18* |
| Neo-adjuvant | Yes  No | 26 | *100* | 6  197 | *3*  *97* |
| Debulking | Optimal  Suboptimal | 24  2 | *92*  *8* | 108  95 | *53*  *47* |
| Series | 19829  20565  26193  9899 | 2  8  2  14 | *8*  *31*  *8*  *54* | 20  38  6  139 | *10*  *19*  *3*  *69* |
| 1 year survival (95% CI) | 100 (NA-100) | | | 96% (83-99) | |
| 3 year survival (95% CI) | 96% (88-100) | | | 63% (56-71) | |
| 5 year survival (95% CI) | 78% (58-100) | | | 36% (28-45) | |
| Median survival (95% CI) | 3392 (1943-NA) | | | 1373 (1220-1541) | |

C. Clinicopathological parameters for subgroups defined by node 3 (Stage)

| Node 4 (TC 166 / 13q12-q14 / Replication & apoptosis) |  | Low (#) | % | High (#) | % |
| --- | --- | --- | --- | --- | --- |
| Age, mean (range) |  | 58 (23-80) | | 61 (39-89) | |
| Taxol treated | Yes  No | 72  15 | *93*  *17* | 94  22 | *81*  *19* |
| Neo-adjuvant | Yes  No | 3  84 | *3.4*  *97* | 3  113 | *3*  *97* |
| Debulking | Optimal  Suboptimal | 40  47 | *46*  *54* | 48  68 | *41*  *59* |
| Series | 19829  20565  26193  9899 | 16  3  68 | *18*  *3*  *78* | 20  22  3  71 | *17*  *19*  *3*  *61* |
| 1 year survival (95% CI) | 98% (94-100) | | | 95% (91-99) | |
| 3 year survival (95% CI) | 53% (42-66) | | | 71% (62-81) | |
| 5 year survival (95% CI) | 19% (10-35) | | | 46% (36-59) | |
| Median survival (95% CI) | 1129 (991-1331) | | | 1662 (1415-2196) | |

D. Clinicopathological parameters for subgroups defined by node 4 (TC 166 / 13q12-q14 / Replication & apoptosis)

| Node 5 (TC 247 / 11q13-q14 / Proliferation & immune response) |  | Low (#) | % | High (#) | % |
| --- | --- | --- | --- | --- | --- |
| Age, mean (range) |  | 59 (39-87) | | 65 (41-89) | |
| Taxol treated | Yes  No | 18  58 | *24*  *76* | 4  36 | *10*  *90* |
| Neo-adjuvant | Yes  No | 2  74 | *2.6*  *97* | 1  39 | *2.5*  *98* |
| Debulking | Optimal  Suboptimal | 41  35 | *54*  *46* | 27  13 | *68*  *33* |
| Series | 19829  20565  26193  9899 | 3  17  3  53 | *3.9*  *22*  *3.9*  *70* | 17  5  18 | *43*  *13*  *45* |
| 1 year survival (95% CI) | 96% (92-100) | | | 92% (84-100) | |
| 3 year survival (95% CI) | 78% (68-88) | | | 57% (42-78) | |
| 5 year survival (95% CI) | 56%% (41-69) | | | 26% (13-52) | |
| Median survival (95% CI) | 2074 (1541-2784) | | | 1220 (1007-2196) | |

E. Clinicopathological parameters for subgroups defined by node 5 (TC 247 / 11q13-q14 / Proliferation & immune response)

| Node 6 (TC 250 / ECM interactions) |  | Low (#) | % | High (#) | % |
| --- | --- | --- | --- | --- | --- |
| Age, mean (range) | 60 (39-87) | | | 60 (48-76) | |
| Taxol treated | Yes  No | 47  12 | *80*  *20* | 11  6 | *65*  *35* |
| Neo-adjuvant | Yes  No | 2  57 | *3.4*  *97* | 17 | *100* |
| Debulking | Optimal  Suboptimal | 29  30 | *49*  *51* | 12  5 | *71*  *29* |
| Series | 19829  20565  26193  9899 | 3  13  3  40 | *5.1*  *22*  *5.1*  *68* | 4  13 | *24*  *77* |
| 1 year survival (95% CI) | 97% (92-100) | | | 94% (84-100) | |
| 3 year survival (95% CI) | 82% (73-94) | | | 62% (41-93) | |
| 5 year survival (95% CI) | 64% (50-80) | | | 33% (14-77) | |
| Median survival (95% CI) | 2267 (1860-NA) | | | 1403 (1088-NA) | |

F. Clinicopathological parameters for subgroups defined by node 6 (TC 250 / ECM interactions)

| Node 7 ( Age) |  | ≤53.7 (#) | % | > 53.7 (#) | % |
| --- | --- | --- | --- | --- | --- |
| Age, mean (range) |  | 48 (39-54) | | 64 (54-87) | |
| Taxol treated | Yes  No | 14  5 | *74*  *26* | 33  7 | *83*  *18* |
| Neo-adjuvant | Yes  No | 1  18 | *5*  *95* | 1  39 | *2.5*  *98* |
| Debulking | Optimal  Suboptimal | 4  15 | *21*  *79* | 25  15 | *63*  *38* |
| Series | 19829  20565  26193  9899 | 9  10 | *47*  *53* | 3  4  3  10 | *7.5*  *10*  *7.5*  *25* |
| 1 year survival (95% CI) | 100 (NA-100) | | | 95% (89-100) | |
| 3 year survival (95% CI) | 77 (59-100) | | | 87 (75-98) | |
| 5 year survival (95% CI) | 44 (24-77) | | | 77 (63-94) | |
| Median survival (95% CI) | 1663 (1415-NA) | | | 2784 (2227-NA) | |

G. Clinicopathological parameters for subgroups defined by node 7 (Age)

| Node 8 (TC 76 / 9p13-p21 / Replication stress) |  | Low (#) | % | High (#) | % |
| --- | --- | --- | --- | --- | --- |
| Age, mean (range) |  | 52 (23-68) | | 60 (35-80) | |
| Taxol treated | Yes  No | 20 | *100* | 15  52 | *22*  *78* |
| Neo-adjuvant | Yes  No | 20 | *100* | 3  64 | *4.5*  *96* |
| Debulking | Optimal  Suboptimal | 11  9 | *55*  *45* | 29  38 | *43*  *57* |
| Series | 20565  26193  9899 | 3  1  16 | *15*  *5*  *80* | 13  2  52 | *19*  *3*  *78* |
| 1 year survival (95% CI) | 100% (NA-100) | | | 97% (93-100) | |
| 3 year survival (95% CI) | 74% (57-97) | | | 46% (34-62) | |
| 5 year survival (95% CI) | 54% (32-90) | | | 11% (4-28) | |
| Median survival (95% CI) | 2284 (1274-NA) | | | 1024 (854-1233) | |

H. Clinicopathological parameters for subgroups defined by node 8 (TC 76 / 9p13-p21 / Replication stress)

| Node 9 (TC 146 / Neurotransmitter signaling) |  | Low (#) | % | High (#) | % |
| --- | --- | --- | --- | --- | --- |
| Age, mean (range) | 57 (35-80) | | | 63 (46-80) | |
| Taxol treated | Yes  No | 32  6 | *84*  *16* | 20  9 | *69*  *31* |
| Neo-adjuvant | Yes  No | 2  36 | *5.3*  *95* | 1  28 | *3.4*  *97* |
| Debulking | Optimal  Suboptimal | 15  23 | *40*  *61* | 14  15 | *48*  *52* |
| Series | 20565  26193  9899 | 7  31 | *18*  *82* | 6  2  21 | *21*  *6.9*  *72* |
| 1 year survival (95% CI) | 100% (NA-100) | | | 93% (84-100) | |
| 3 year survival (95% CI) | 62% (46-83) | | | 26% (13-53) | |
| 5 year survival (95% CI) | 14% (4-44) | | | 7% (1-41) | |
| Median survival (95% CI) | 1159 (1034-1392) | | | 735 (641-1159) | |

I. Clinicopathological parameters for subgroups defined by node 9 (TC 146 / Neurotransmitter signaling)
