## Supplementary figures tables and methods for "Transcriptional pattern enriched for synaptic signaling is associated with shorter survival of patients with high-grade serous ovarian cancer": Supplementary Table S6.docx

**Supplementary Table S6.** Clinicopathological parameters per survival tree node in advanced-stage platinum-treated high-grade serous ovarian cancer.

| Node 1 (TC 166 / 13q12-q14 / Replication & apoptosis) |  | Low (#) | % | High (#) | % |
| --- | --- | --- | --- | --- | --- |
| Stage | 3  4 | 99  13 | *88*  *12* | 133  20 | *87*  *13* |
| Age, mean (range) |  | 59 (23-80) | | 61 (38-89) | |
| Taxol treated | Yes  No | 90  22 | *80*  *20* | 120  33 | *78*  *22* |
| Neo-adjuvant | Yes  No | 5  107 | *4.5*  *96* | 7  146 | *4.6*  *95* |
| Debulking | Optimal  Suboptimal | 46  66 | *41*  *59* | 86  67 | *56*  *44* |
| Series | 19829  20565  26193  9899 | 19  5  88 | *17*  *4.5*  *79* | 25  32  4  92 | *16*  *21*  *2.6*  *60* |
| 1 year survival (95% CI) | 94% (89-98) | | | 95% (91-98) | |
| 3 year survival (95% CI) | 44% (34-55) | | | 63% (55-72) | |
| 5 year survival (95% CI) | 15% (8-29) | | | 41% (32-52) | |
| Median survival | 991 (823-1159) | | | 1464 (1281-1861) | |

A. Clinicopathological parameters for subgroups defined by node 1 (TC 166 / 13q12-q14 / Replication & apoptosis)

| Node 2 (TC 121 / Neuronal development) |  | Low (#) | % | High (#) | % |
| --- | --- | --- | --- | --- | --- |
| Stage | 3  4 | 76  11 | *87*  *13* | 23  2 | *92*  *8* |
| Age, mean (range) |  | 58 (23-80) | | 61 (33-80) | |
| Taxol treated | Yes  No | 71  16 | *82*  *18* | 19  6 | *76*  *6* |
| Neo-adjuvant | Yes  No | 5  82 | *5.7*  *94* | 25 | *100* |
| Debulking | Optimal  Suboptimal | 36  51 | *41*  *59* | 10  15 | *40*  *60* |
| Series | 20565  26193  9899 | 15  5  67 | *17*  *5.7*  *77* | 4  21 | *16*  *84* |
| 1 year survival (95% CI) | 98% (94-100) | | | 80% (66-97) | |
| 3 year survival (95% CI) | 51% (40-64) | | | 18% (7-48) | |
| 5 year survival (95% CI) | 18% (10-34) | | | 9% (2-49) | |
| Median survival (95% CI) | 1112 (988-1312) | | | 671 (549-854) | |

B. Clinicopathological parameters for subgroups defined by node 2 (TC 121 / Neuronal development)

| Node 3 (TC 247 / 11q13-q14 / Proliferation & immune response) |  | Low (#) | % | High (#) | % |
| --- | --- | --- | --- | --- | --- |
| Stage | 3  4 | 22 | *100* | 54  11 | *83*  *17* |
| Age, mean (range) |  | 55 (35-76) | | 59 (23-80) | |
| Taxol treated | Yes  No | 20  2 | *91*  *9* | 51  14 | *79*  *22* |
| Neo-adjuvant | Yes  No | 1  21 | *4.5*  *96* | 4  61 | *6.2*  *94* |
| Debulking | Optimal  Suboptimal | 12  10 | *55*  *46* | 24  41 | *37*  *63* |
| Series | 20565  26193  9899 | 7  1  14 | *32*  *4.5*  *64* | 8  4  53 | *12*  *6.2*  *82* |
| 1 year survival (95% CI) | 95% (87-100) | | | 98% (95-100) | |
| 3 year survival (95% CI) | 70% (53-94) | | | 46% (34-62) | |
| 5 year survival (95% CI) | 39% (21-72) | | | 7% (1-34) | |
| Median survival (95% CI) | 1342 (1112-NA) | | | 1007 (796-1233) | |

C. Clinicopathological parameters for subgroups defined by node 3 (TC 247 / 11q13-q14 / Proliferation & immune response)

| Node 4 (TC 76 / 9p13-p21 / Replication stress) |  | Low (#) | % | High (#) | % |
| --- | --- | --- | --- | --- | --- |
| Stage | 3  4 | 16  2 | *89*  *11* | 38  9 | *81*  *19* |
| Age, mean (range) |  | 52 (23-70) | | 62 (39-80) | |
| Taxol treated | Yes  No | 17  1 | *94*  *5.6* | 34  13 | *72*  *28* |
| Neo-adjuvant | Yes  No | 1  17 | *5.6*  *94* | 3  44 | *6.4*  *94* |
| Debulking | Optimal  Suboptimal | 7  11 | *39*  *61* | 17  30 | *36*  *64* |
| Series | 20565  26193  9899 | 1  17 | *5.6*  *94* | 7  4  36 | *15*  *8.5*  *77* |
| 1 year survival (95% CI) | 94% (84-100) | | | 100% (NA-100) | |
| 3 year survival (95% CI) | 58% (37-92) | | | 41% (28-61) | |
| 5 year survival (95% CI) | 29% (7-100) | | | 4% (2-28) | |
| Median survival (95% CI) | 1739 (843-NA) | | | 937 (793-1159) | |

D. Clinicopathological parameters for subgroups defined by node 8 (TC 76 / 9p13-p21 / Replication stress)

| Node 5 (TC 247 / 11q13-q14 / Proliferation & immune response) |  | Low (#) | | % | High (#) | % |
| --- | --- | --- | --- | --- | --- | --- |
| Stage | 3  4 | 28  72 | | *28*  *72* | 12  41 | *23*  *77* |
| Age, mean (range) |  | 59 (38-87) | | | 64 (41-87) | |
| Taxol treated | Yes  No | 75  25 | *75*  *25* | | 45  8 | *85*  *15* |
| Neo-adjuvant | Yes  No | 6  94 | *6*  *94* | | 1  52 | *1.9*  *98* |
| Debulking | Optimal  Suboptimal | 53  47 | *53*  *47* | | 33  20 | *62*  *38* |
| Series | 19829  20565  26193  9899 | 4  25  4  67 | *4*  *25*  *4*  *67* | | 21  7  25 | *40*  *13*  *47* |
| 1 year survival (95% CI) | 96% (92-100) | | | | 92% (85-100) | |
| 3 year survival (95% CI) | 70% (61-80) | | | | 50% (37-68) | |
| 5 year survival (95% CI) | 50% (39-63) | | | | 21% (11-43) | |
| Median survival (95% CI) | 1769 (1454-2227) | | | | 1037 (859-1464) | |

E. Clinicopathological parameters for subgroups defined by node 5 (TC 247 / 11q13/q14 / Proliferation & immune response)

| Node 6 (TC 166 / 13q12-q14 / Replication & apoptosis) |  | Low (#) | % | High (#) | % |
| --- | --- | --- | --- | --- | --- |
| Stage | 3  4 | 18  3 | *86*  *14* | 30  2 | *94*  *6* |
| Age, mean (range) |  | 60 (47-76) | | 66 (41-89) | |
| Taxol treated | Yes  No | 20  1 | *95*  *4.8* | 25  7 | *78*  *22* |
| Neo-adjuvant | Yes  No | 1  20 | *4.8*  *95* | 32 | *100* |
| Debulking | Optimal  Suboptimal | 16  5 | *76*  *24* | 17  15 | *53*  *47* |
| Series | 19829  20565  9899 | 10  1  10 | *48*  *4.8*  *48* | 11  6  15 | *34*  *19*  *47* |
| 1 year survival (95% CI) | 100 (NA-100) | | | 87% (76-100) | |
| 3 year survival (95% CI) | 84% (65-100) | | | 30% (16-54) | |
| 5 year survival (95% CI) | 37% (17-81) | | | 12% (3-41) | |
| Median survival (95% CI) | 1373 (1220-NA) | | | 763 (549-1464) | |

F. Clinicopathological parameters for subgroups defined by node 6 (TC 166 / 13q12-q14 / Replication & apoptosis)

| Node 7 (TC 250 / ECM interactions) |  | Low (#) | % | High (#) | % |
| --- | --- | --- | --- | --- | --- |
| Stage | 3  4 | 64  10 | *87*  *14* | 21  5 | *81*  *19* |
| Age, mean (range) |  | 58 (38-87) | | 61 (44-78) | |
| Taxol treated | Yes  No | 58  16 | *78*  *22* | 17  9 | *65*  *35* |
| Neo-adjuvant | Yes  No | 5  69 | *6.8*  *93* | 1  25 | *3.8*  *96* |
| Debulking | Optimal  Suboptimal | 37  37 | *50*  *50* | 16  10 | *62*  *39* |
| Series | 19829  20565  26193  9899 | 4  19  3  48 | *5.4*  *26*  *4.1*  *65* | 6  1  19 | *23*  *3.8*  *73* |
| 1 year survival (95% CI) | 97% (94-100) | | | 92% (83-100) | |
| 3 year survival (95% CI) | 75% (65-87) | | | 64% (48-86) | |
| 5 year survival (95% CI) | 58% (45-73) | | | 28% (13-63) | |
| Median survival (95% CI) | 1983 (1663-2685) | | | 1159 (970-NA) | |

G. Clinicopathological parameters for subgroups defined by node 7 (TC 250 / ECM interactions)

| Node 8 (Age) |  | ≤ 58 (#) | % | > 58 (#) | % |
| --- | --- | --- | --- | --- | --- |
| Stage | 3  4 | 33  7 | *83*  *18* | 31  3 | *91*  *8.8* |
| Age, mean (range) |  | 51 (38-58) | | 67 (58-87) | |
| Taxol treated | Yes  No | 30  10 | *75*  *25* | 28  6 | *82*  *18* |
| Neo-adjuvant | Yes  No | 3  37 | *7.5*  *93* | 2  32 | *5.9*  *94* |
| Debulking | Optimal  Suboptimal | 15  25 | *38*  *63* | 22  12 | *65*  *35* |
| Series | 19829  20565  26193  9899 | 1  13  1  25 | *2.5*  *33*  *2.5*  *63* | 3  6  2  23 | *8.8*  *18*  *5.9*  *68* |
| 1 year survival (95% CI) | 98% (93-100) | | | 97% (92-100) | |
| 3 year survival (95% CI) | 69% (56-86) | | | 88% (77-100) | |
| 5 year survival (95% CI) | 47% (32-69) | | | 73% (57-93) | |
| Median survival (95% CI) | 1687 (1415-2231) | | | 3233 (2074-NA) | |

H. Clinicopathological parameters for subgroups defined by node 8 (Age)
