## Supplementary figures tables and methods for "Transcriptional pattern enriched for synaptic signaling is associated with shorter survival of patients with high-grade serous ovarian cancer": Supplementary Table S9.docx

**Supplementary Table S8.** Expression and function of the top 20 genes in TC 121

| **Symbol** | **Common name** | **Tissue expression & function** |
| --- | --- | --- |
| LHX8 | LIM homeobox 8 | Essential transcription factor for the development of neurons, teeth and oocytes. Appears to be regulated by methylation.^22^ LHX8 methylation is an important marker for precancerous cervical lesions.^23^ |
| DMRTC2 | DMRT like family C2 | Related to male and female germ cell development, downstream of cyclin E.^24,25^ |
| EN1 | engrailed homeobox 1 | Plays a role in central nervous system development and is hypermethylated in HGSOC, colorectal, prostate and breast cancer.^26,27^ |
| ESC1 | ESX homeobox 1 | X-linked, germ cell-specific gene. It is related to cell cycle progression and transcription during spermatogenesis.^28^ Suppresses KRAS signaling in xenograft models, resulting in growth inhibition.^29^ |
| SOX30 | SRY-box 30 | Epigenetically regulated transcription factor overexpressed in oocytes.^30^ Related to better survival of advanced lung, bladder, and ovarian cancer.^31–33^ Activates p53-signaling.^34^ |
| FOXE3 | forkhead box E3 | Methylation is related to survival of clear cell renal cell carcinoma patients.^35^ |
| CARTPT | CART prepropeptide | Methylation-regulated inhibitor of follicle development.^36^ |
| BEND4 | BEN domain containing 4 | Hypermethylated in oropharyngeal , colorectal, and lung cancer.^37–39^ |
| SOGA3 | SOGA family member 3 | - |
| ZNF536 | zinc finger protein 536 | Induces neuronal differentiation.^40^ |
| TMEM151B | transmembrane protein 151B | - |
| TAF7L | TATA-box binding protein associated factor 7 like | Germ cell-specific and plays a role in spermatogenesis^41^ |
| OTP | orthopedia homeobox | Key player in hypothalamus development. Expression is related to better OS of pulmonary carcinoid patients.^42^ |
| ZIC3 | Zic family member 3 | Upregulated by EN1 in breast cancer cells.^43^ Negatively regulated by miR-564, resulting in proliferation and motility.^44^ |
| DRAXIN | dorsal inhibitory axon guidance protein | Important for neuron development, inhibits axonal growth through Akt.^45^ Potentially plays a role in proliferation and apoptosis of lung cancer cells.^46^ |
| LRRC14B | leucine rich repeat containing 14B | - |
| ELAVL4 | ELAV like RNA binding protein 4 | Marker of neuronal phenotype and essential for neuronal development.^47^ It is highly immunogenic, possibly related to SCLC-autoimmunity and proliferation of NSCLC.^48,49^ |
| GSC | goosecoid homeobox | Transcription factor related to neural crest development. Increased in ovarian cancer stem-like cells and related to poor drug response and OS.^50,51^ |
| CCDC144NL-AS1 | CCDC144NL antisense RNA 1 | Related to OS of BRCA-wild type ovarian and colorectal cancer.^52,53^ In endometriosis cell line models related to vimentin expression and migration.^54^ Prevents conversion of pluripotent stem cells back to naïve-like state.^55^ |
| LECT1 | leukocyte cell derived chemotaxin 1 | Angiogenesis inhibitor, increases chondrocyte growth.^56^ |
