## Supplementary figures and images for "Transcriptional pattern enriched for synaptic signaling is associated with shorter survival of patients with high-grade serous ovarian cancer"

### Supplementary figure S1.pdf

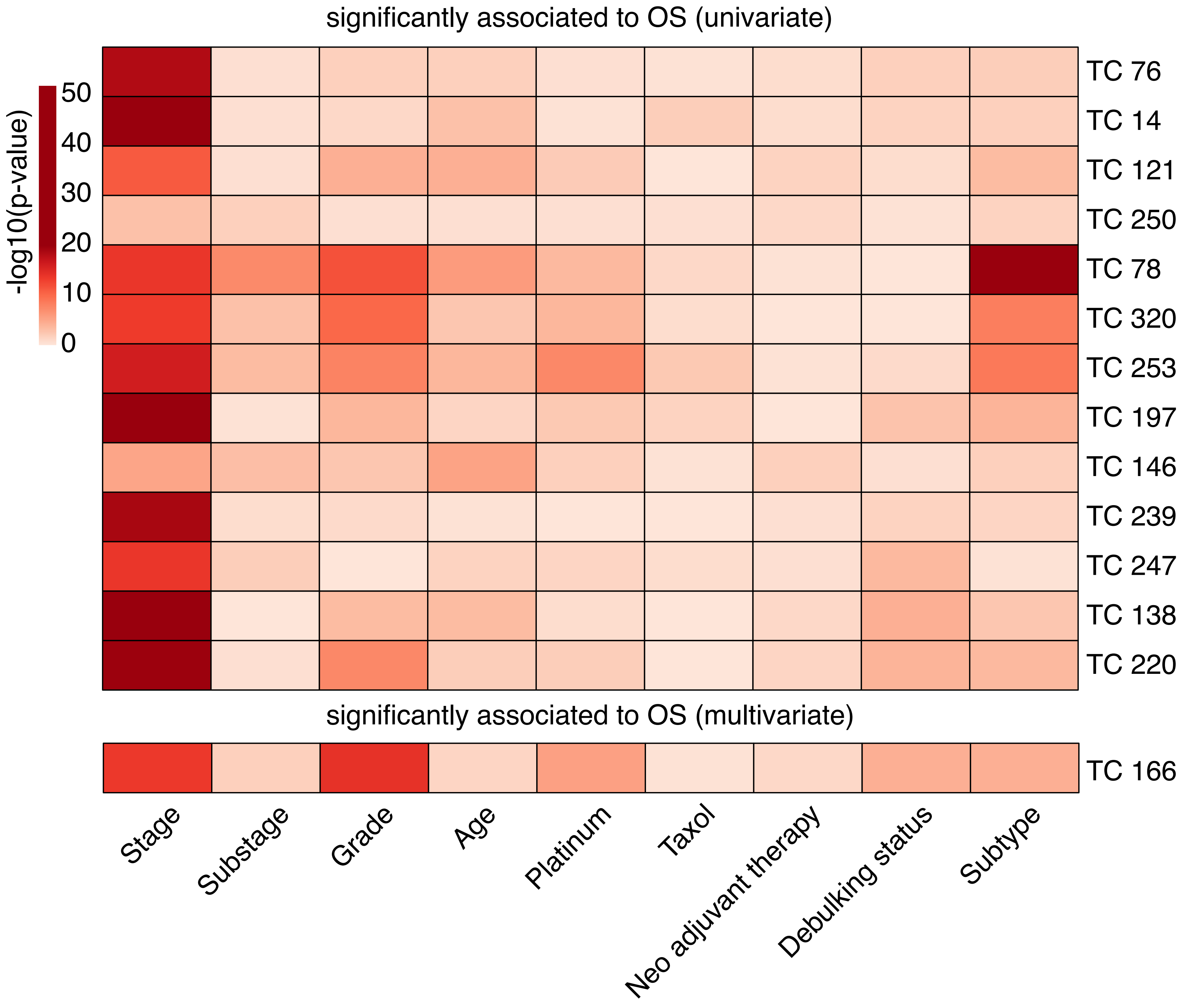

### Supplementary figure S2.pdf

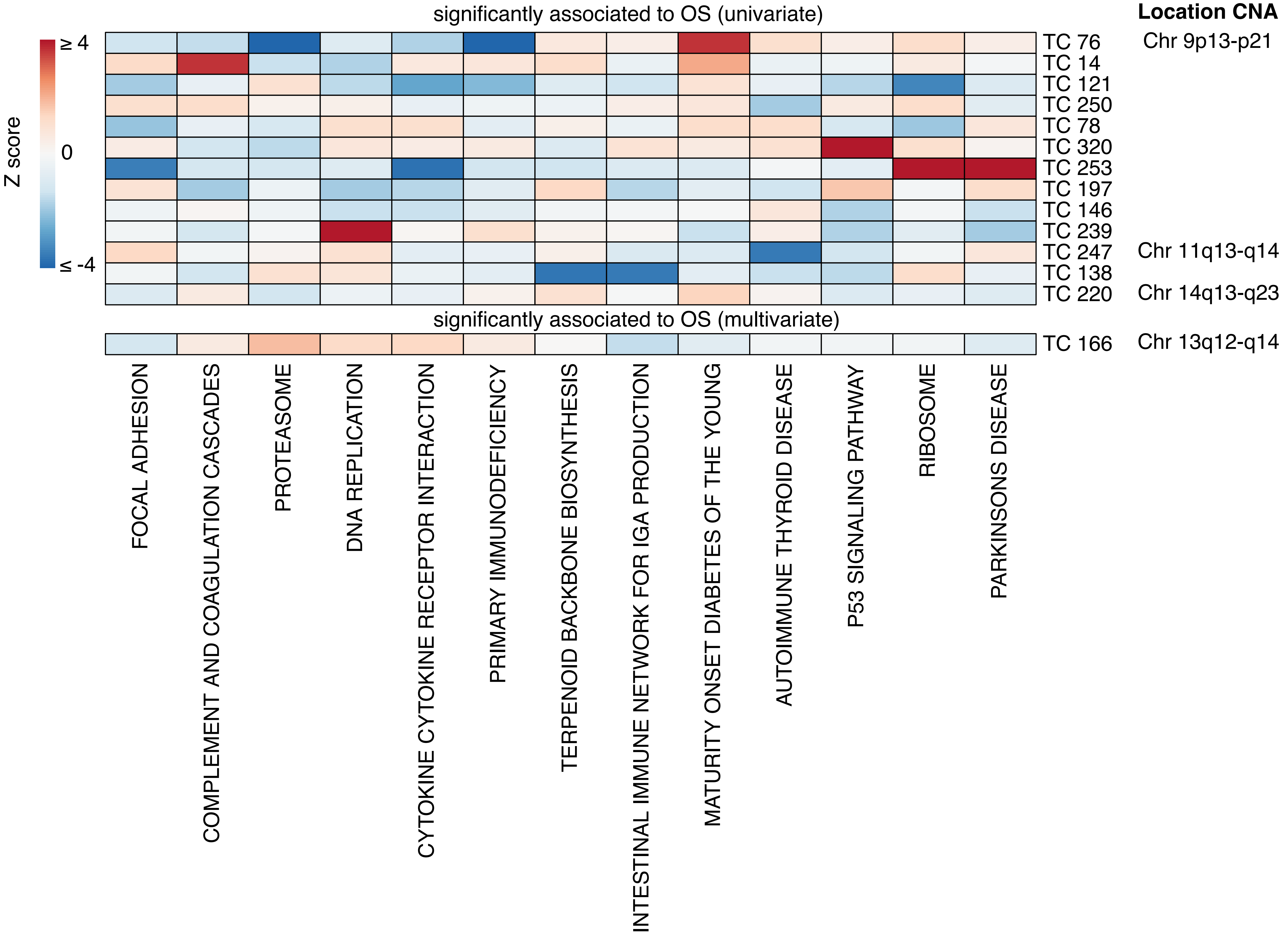

### Supplementary figure S3.pdf

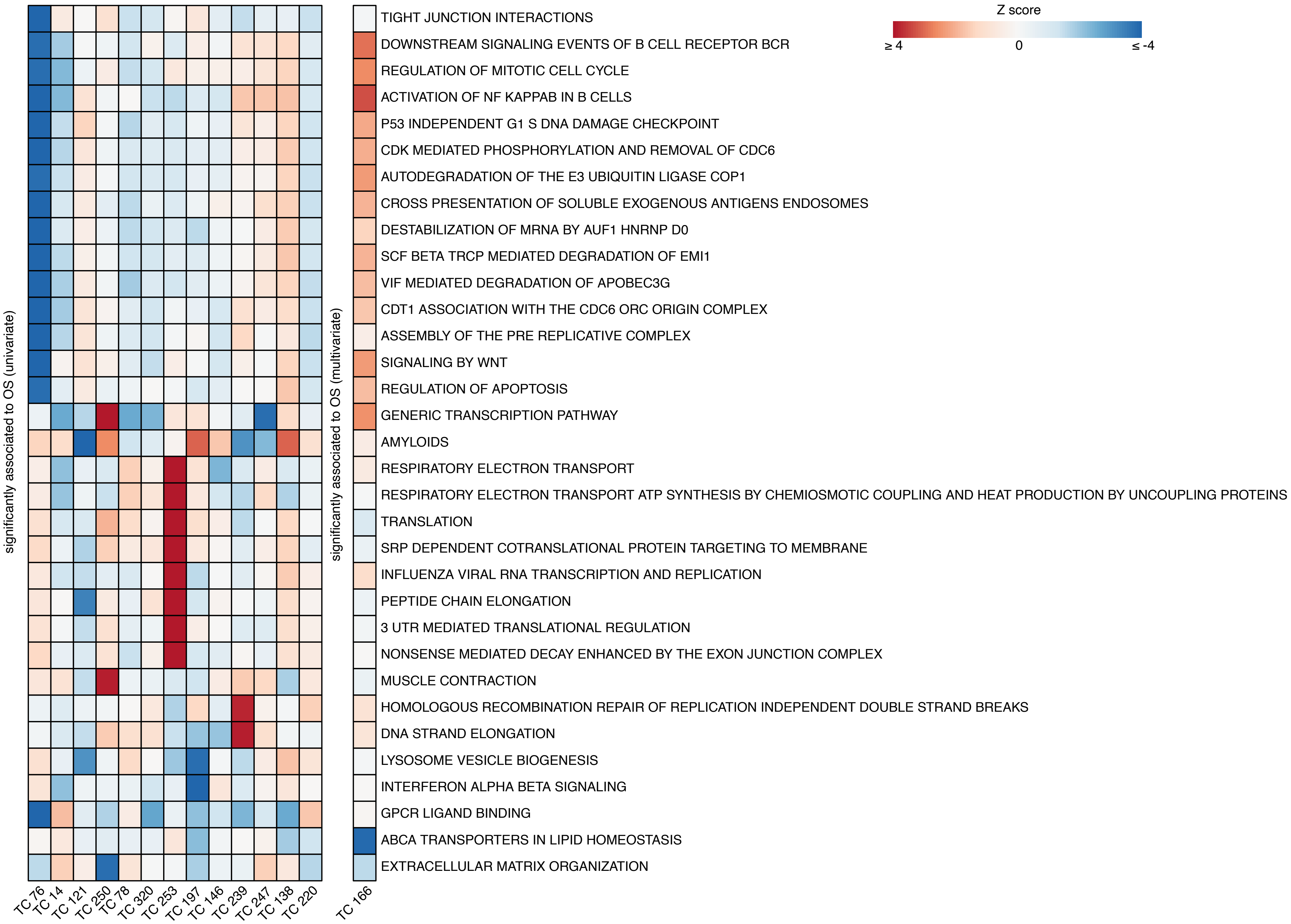

### Supplementary figure S4.pdf

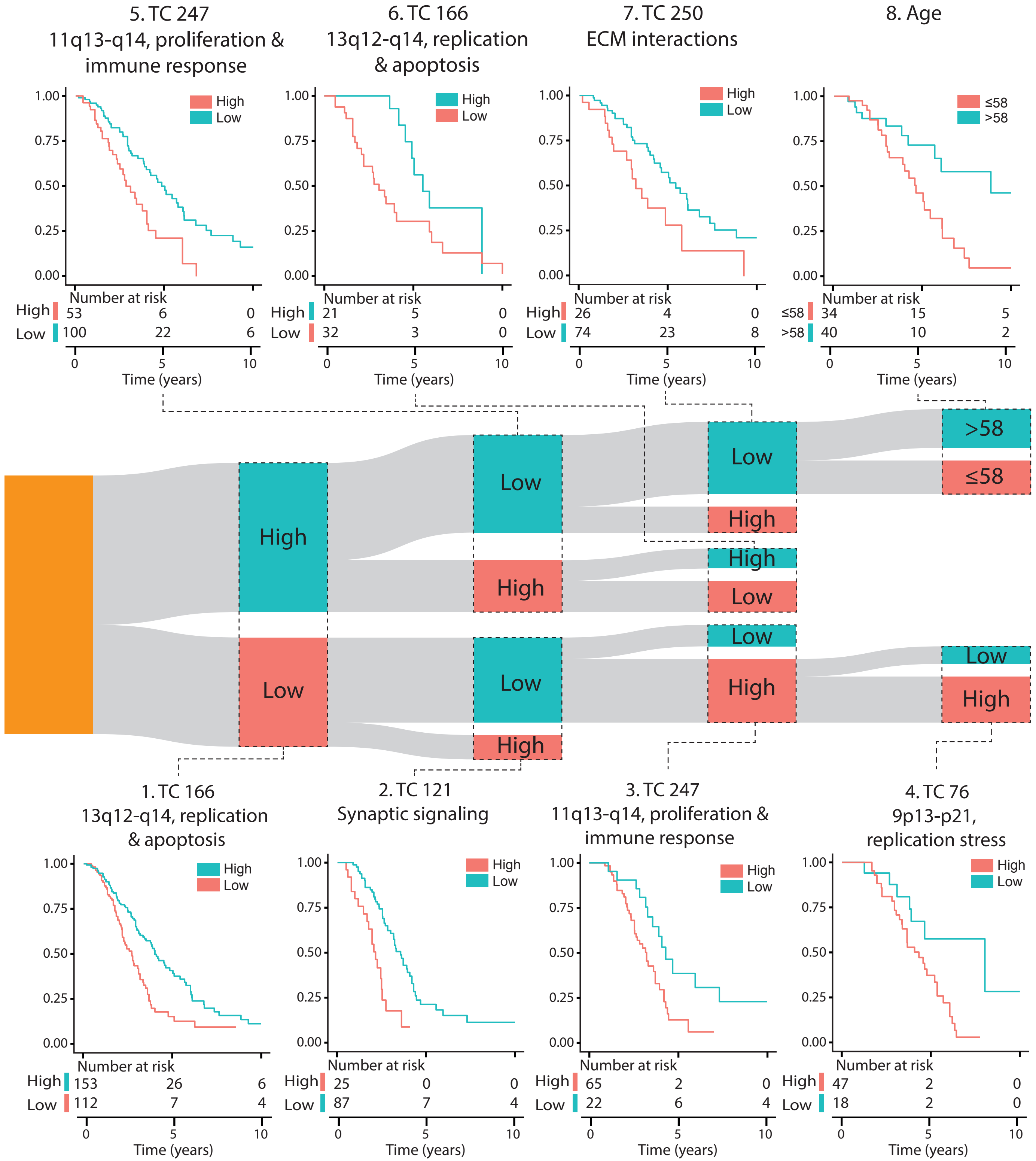

### Supplementary figure S5.pdf

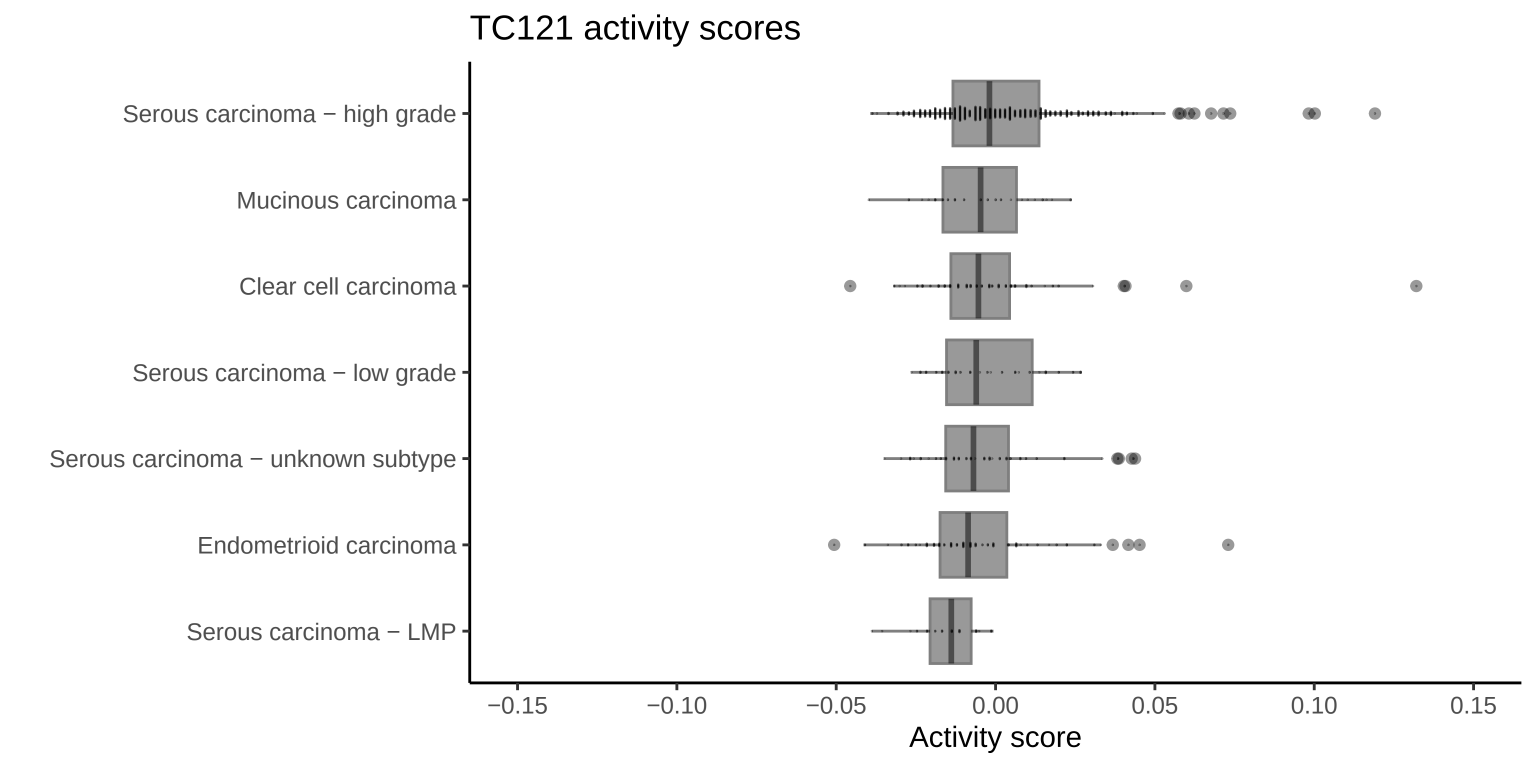

### Supplementary figure S6.pdf

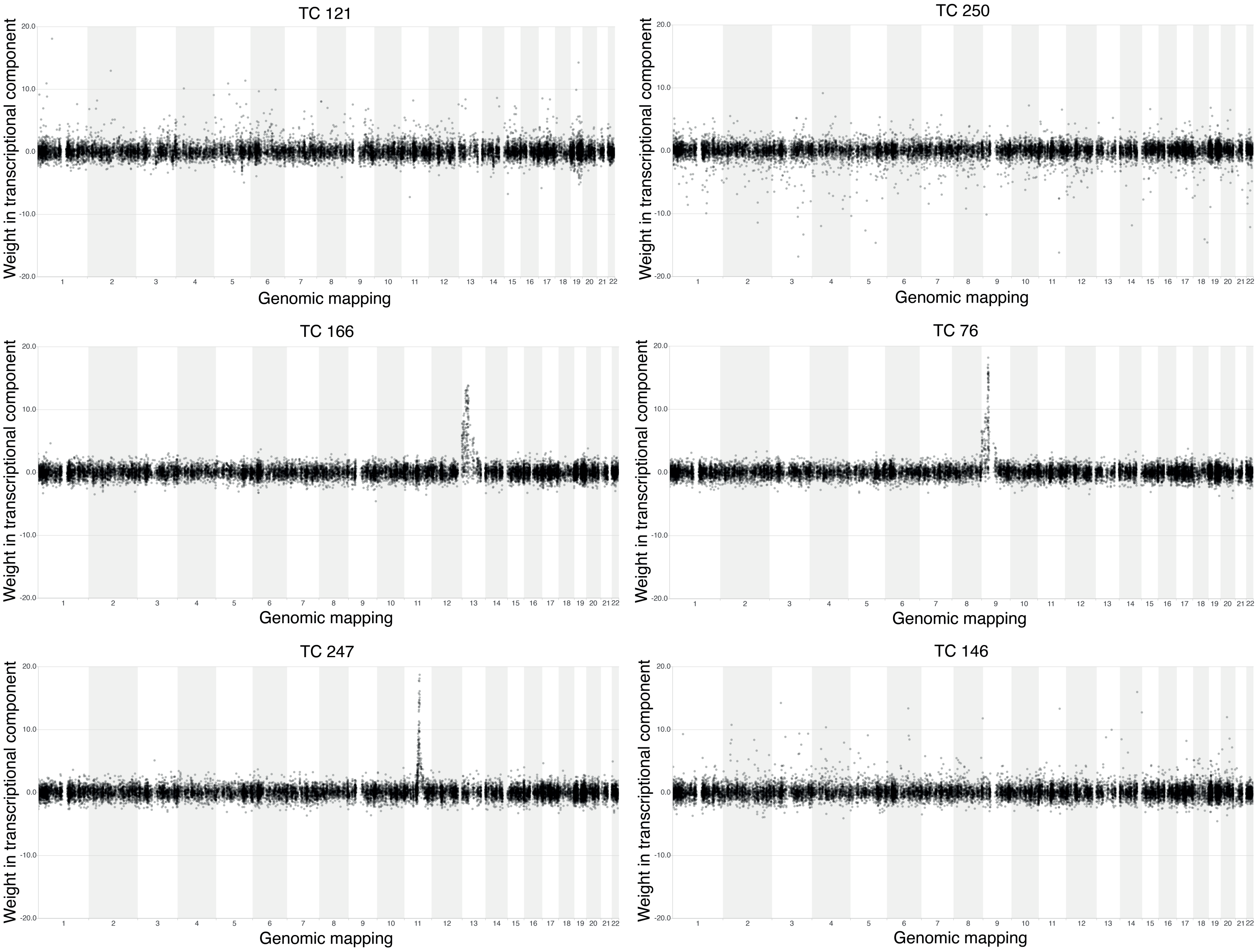

### Supplementary figure S7.pdf

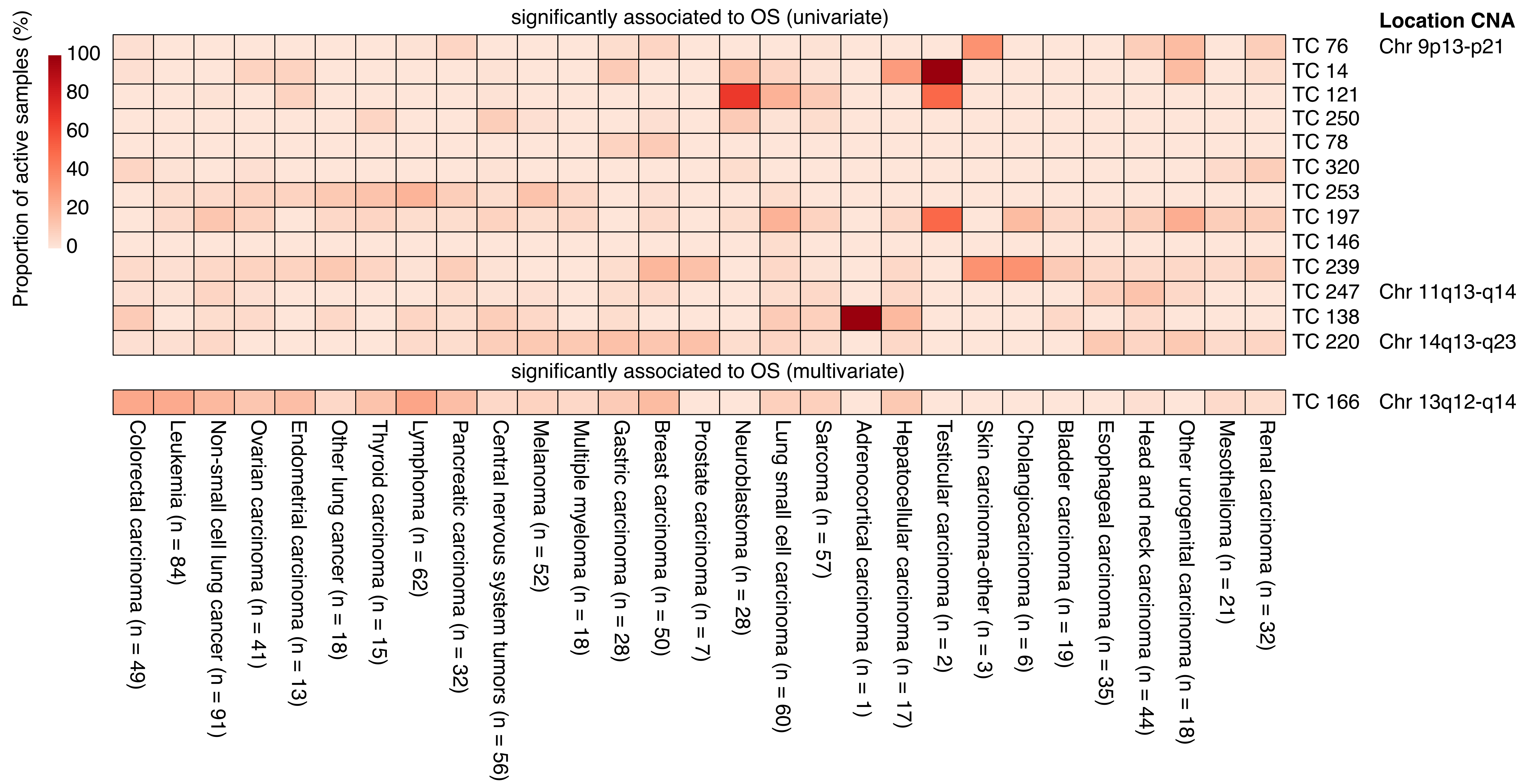

### Supplementary figure S8.pdf

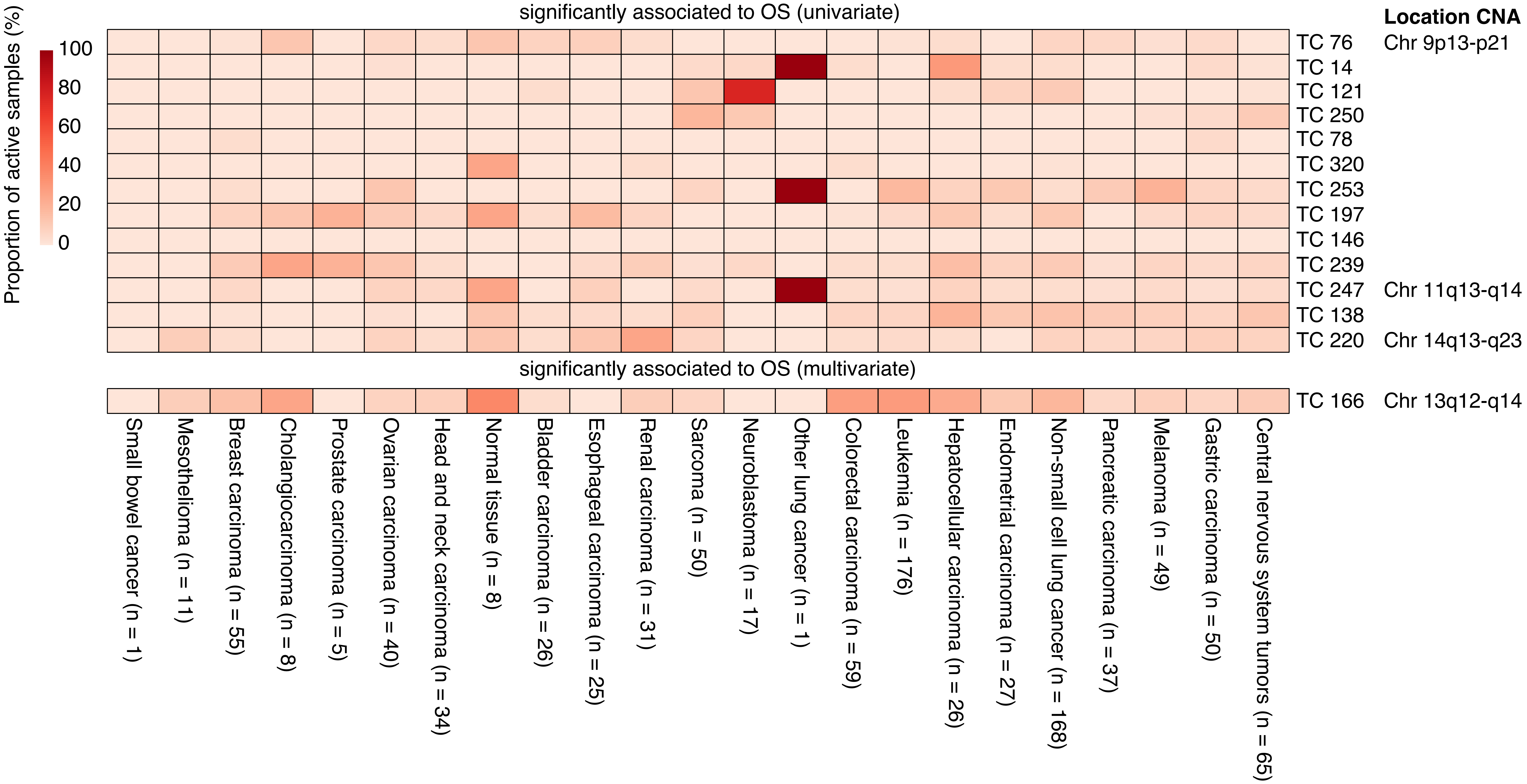

### Supplementary figure S9.pdf

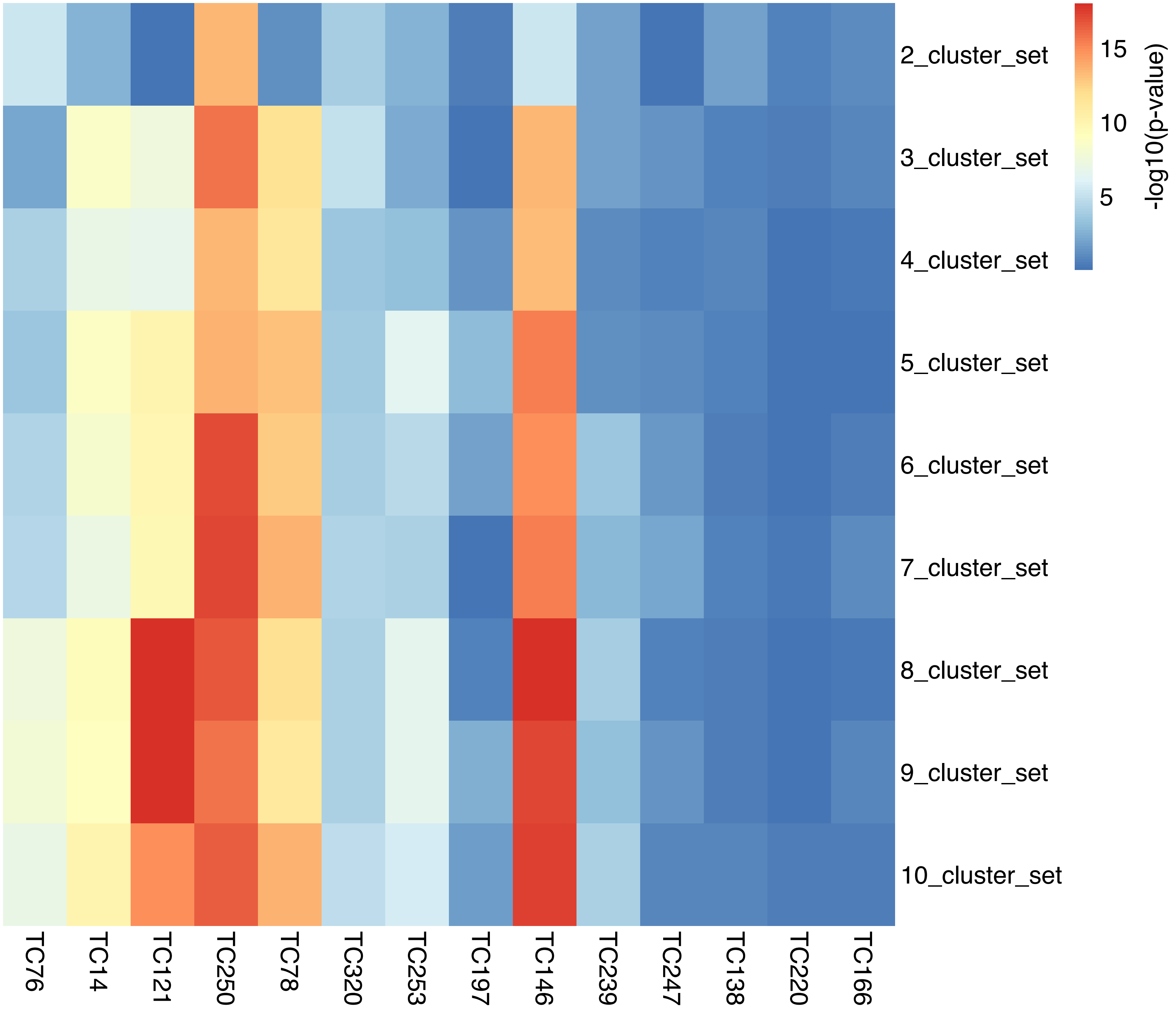

### Supplementary figure S10.png

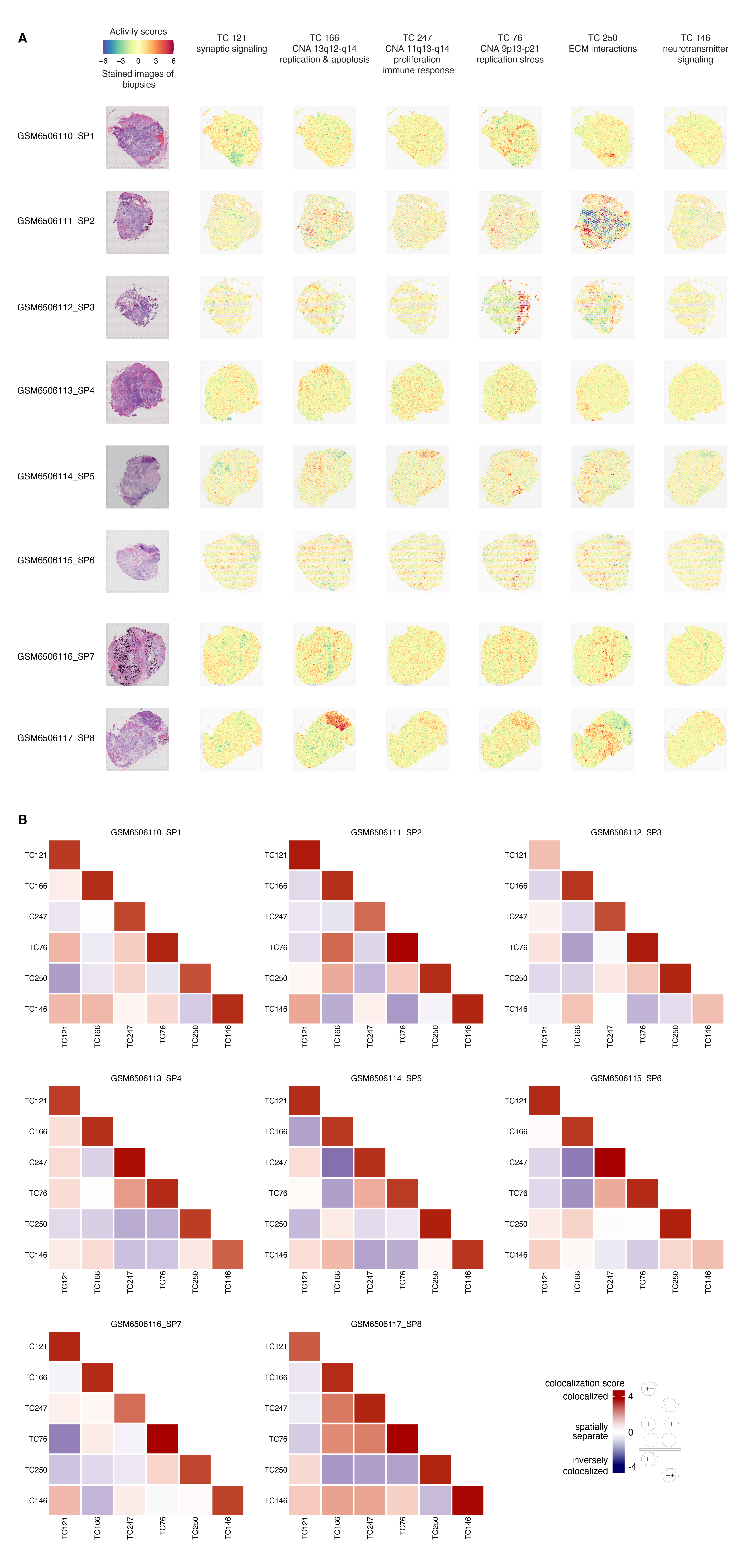

### Supplementary figure S11.pdf

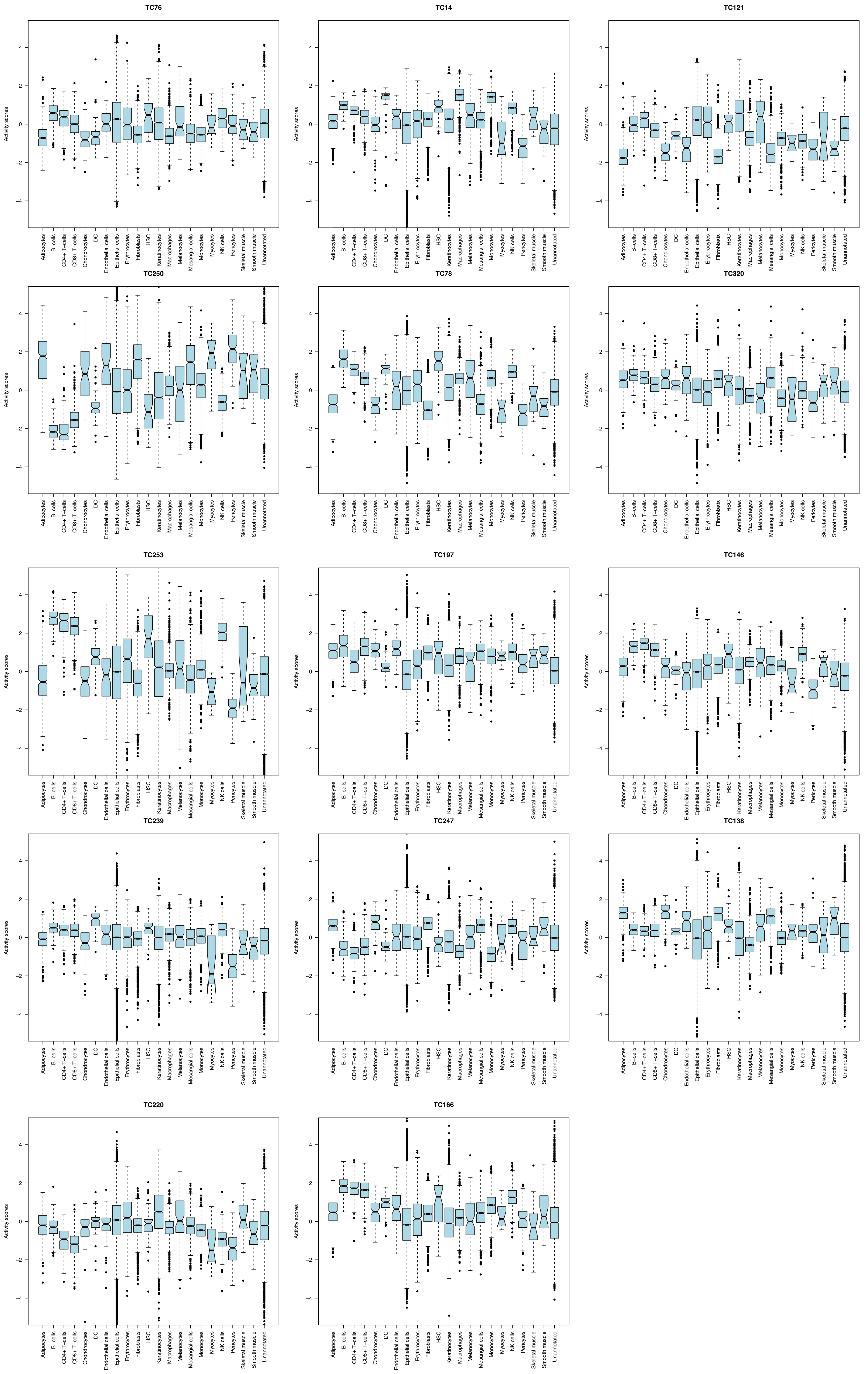
